# Transcription-driven Chromatin Repression of Intragenic Promoters

**DOI:** 10.1101/279414

**Authors:** Mathias Nielsen, Ryan Ard, Xueyuan Leng, Maxim Ivanov, Peter Kindgren, Vicente Pelechano, Sebastian Marquardt

## Abstract

Progression of RNA polymerase II (RNAPII) transcription relies on the appropriately positioned activities of elongation factors. The resulting profile of factors and chromatin signatures along transcription units provides a “positional information system” for transcribing RNAPII. Here, we investigate a chromatin-based mechanism that suppresses intragenic initiation of RNAPII transcription. We demonstrate that RNAPII transcription across gene promoters represses their function in plants. This repression is characterized by reduced promoter-specific molecular signatures and increased molecular signatures associated with RNAPII elongation. The FACT histone chaperone complex is required for this repression mechanism. Genome-wide mapping of Transcription Start Sites (TSSs) reveals thousands of discrete intragenic TSS positions in FACT mutants. Histone 3 lysine 4 mono-methylation poises exonic sites to initiate RNAPII transcription in FACT mutants. Uncovering the mechanism for intragenic TSS repression through the act of RNAPII elongation has important implications for understanding pervasive RNAPII transcription and the regulation of transcript isoform diversity.

## Introduction

Plasticity at the beginning and end of transcripts multiplies the RNA species that can be generated from genomes. RNA results from RNAPII activity at genes, but also from pervasive transcription of abundant non-coding genomic regions (Jensen et al., 2013; Mellor et al., 2016). Pervasive transcription may result in overlapping transcripts, for example by initiating intragenic transcription leading to the production of alternative transcript isoforms (Davuluri et al., 2008). Alternative Transcription Start Sites (TSSs) expand RNA isoform diversity, may result in functionally different RNA and proteins specific to disease, and allow for multiple transcriptional inputs from a single gene (Arner et al., 2015; Wiesner et al., 2015). However, the mechanisms of alternative TSS activation, repression and regulation are poorly understood in higher eukaryotes.

Repression of a gene promoter by overlapping RNAPII transcription was originally described for two tandemly arranged human α-globin gene copies (Proudfoot, 1986). Read-through transcription from the upstream α-globin gene positions the downstream promoter in the middle of transcription unit spanning both gene copies. The repression of the downstream promoter through the act of RNAPII transcription is referred to as Transcriptional Interference (TI) (Ard et al., 2017). The core of this mechanism relies on the progression of RNAPII transcription through distinct stages (Buratowski, 2009). Each stage is characterized by the co-transcriptional recruitment of factors involved in nascent RNA processing and chromatin modifications (Li et al., 2007). Dynamic phosphorylation of residues in the C-terminal YSPTSPS repeat region of the largest RNAPII subunit coordinates the progression through transcription cycle by recruiting stage-specific factors (Corden, 2013; Eick and Geyer, 2013). Metagene analyses of stage-specific transcription factors and chromatin signatures in diverse organisms strikingly visualizes many common changes associated with RNAPII progression from the beginning to the end of active transcription units (Descostes et al., 2014; Gerstein et al., 2010; Hajheidari et al., 2013; Kharchenko et al., 2011; Mayer et al., 2010; Pokholok et al., 2005). These signatures provide a “positional information system” (POINS) for RNAPII to coordinate molecular events required for each stage of transcription (Buratowski, 2009).

An important functional outcome of co-transcriptional chromatin changes involves the suppression of cryptic intragenic TSSs. Histone 3 lysine 36 methylation (H3K36me) is characteristic of RNAPII elongation in many organisms (Bannister et al., 2005; Bell et al., 2007; Guenther et al., 2007; Mahrez et al., 2016). H3K36me prevents RNAPII transcription initiation from cryptic promoters within gene bodies by mediating histone de-acetylation in yeast (Carrozza et al., 2005; Keogh et al., 2005; Venkatesh et al., 2012). Chromatin-based repression of intragenic promoters is tightly linked to the activity of histone chaperones (Cheung et al., 2008; Kaplan et al., 2003). The FACT (FAcilitates Chromatin Transcription) complex, consisting of SSRP1 and SPT16, contributes to this activity across taxa (Belotserkovskaya et al., 2003; Orphanides et al., 1999). SPT16 was initially characterized as SPT (*suppressor of Ty*) gene that is required for the suppression of gene promoters by read-through transcription initiating from adjacent upstream Ty or δ-element insertion (Clark-Adams and Winston, 1987; Malone et al., 1991). RNAPII read-through transcription of upstream genes due to inefficient termination can elicit suppression of downstream gene promoters by TI (Ard et al., 2017; Porrua and Libri, 2015; Proudfoot, 2016). Transcripts overlapping gene promoters may also arise from pervasive RNAPII transcription of long non-coding RNA (lncRNA) and suppress initiation by FACT-dependent TI (Ard and Allshire, 2016; Hainer et al., 2012). In mammals, a combination of FACT, H3K36me, and gene-body DNA methylation suppresses intragenic initiation (Carvalho et al., 2013; Neri et al., 2017). Co-transcriptional chromatin signatures are common across species, yet their roles in the regulation of intragenic TSSs often awaits experimental validation.

Many factors characterizing POINS are active in plants (Hajheidari et al., 2013; Van Lijsebettens and Grasser, 2014). The *Arabidopsis* FACT complex is physically associated with multiple RNAPII elongation factors, chromatin modifiers and elongation specific RNAPII isoforms (Antosz et al., 2017; Duroux et al., 2004). Reduced FACT activity results in developmental defects (Lolas et al., 2010) linked to imprinting (Ikeda et al., 2011), yet the role of FACT in TSS selection in plants is unclear. Genome-wide TSS mapping in *Arabidopsis* suggest that a choice between alternative TSSs exists for most transcripts (Tokizawa et al., 2017). Protein isoform diversity control in response to light through regulated of TSS choice underpins the biological significance of this mechanism (Ushijima et al., 2017). TSS choice may also regulate gene expression at the level of translation by the inclusion of an upstream open reading frame (uORF) (von Arnim et al., 2014). Despite the functional significance of alternative TSS choice little is known about the molecular mechanisms regulating this phenomenon in plants.

Here, we demonstrate the repressive effect of RNAPII elongation across gene promoters in *Arabidopsis*. We identify chromatin and RNAPII signatures associated with this form of gene regulation by “repressive transcription” that relies on chromatin remodeling by the FACT complex. We identify several thousand FACT-sensitive intragenic TSSs, revealing a role for FACT in preventing initiation of RNAPII transcription from within transcription units. Our study characterizes the molecular events involved in repressing RNAPII initiation by the process of RNAPII elongation for the first time in the context of a multicellular organism.

## Results

### Gene promoter repression by upstream RNAPII transcription in Arabidopsis thaliana

To investigate gene repression through the act of RNAPII transcription across promoter regions in higher organisms, we performed a literature screen of *Arabidopsis* T-DNA insertion mutants with loss-of-function phenotypes (Lloyd and Meinke, 2012). This specific type of T-DNA mutants must: 1.) be inserted upstream of gene TSSs, 2.) show read-through transcription into downstream genes, and 3.) segregate as recessive loss-of-function phenotype. Application of these criteria identified the *quasimodo1-1* (*qua1-1*) and *red fluorescence in darkness 1-1* (*rfd1-1*) mutants as candidate mutants for further analysis (Bouton et al., 2002; Hedtke and Grimm, 2009).

*QUASIMODO1* (*QUA1*) encodes a glycosyltransferase required for the biosynthesis of cell-adhesion promoting pectins (Bouton et al., 2002). The *qua1-1* T-DNA mutation is inserted 117 bp upstream of the annotated translational start site (Fig. 1A; SFig. 1A). The cell-adhesion defect in *qua1-1* results in dwarfed growth and ruthenium red staining of dark grown *qua1-1* hypocotyls (Fig. 1B). We detect elevated *QUA1* expression in *qua1-1* compared to wild type by RT-qPCR (Fig. 1C). Northern blotting reveals an abundant *T-DNA-QUA1* compound transcript in *qua1-1* instead of the *QUA1* mRNA (Fig. 1D) (Bouton et al., 2002). The extended transcript detected in *qua1-1* corresponds to a predicted transcript initiating within the T-DNA and extending into the downstream *QUA1* gene (SFig. 1B). *RED FLUORESCENT IN DARKNESS 1 (RFD1)* encodes RibA1, the first enzyme in the plant riboflavin biosynthesis pathway (Hedtke and Grimm, 2009). The T-DNA insertion is located 307 bp upstream of the *RFD1* translation start (Fig. 1E; SFig. 1C). Under standard light conditions, soil-grown homozygous *rfd1-1* mutants die as white cotyledons (Fig. 1F) (Hedtke and Grimm, 2009). However, we are able to grow homozygous *rfd1-1* mutants to seed under low light conditions, enabling comparative analysis of the *RFD1* transcript pattern in wild type and homozygous *rfd1-1* mutants. Although RT-qPCR analysis shows about 20-times higher *RFD1* expression in *rfd1-1* compared to wild type (Fig. 1G), northern blotting reveals an abundant *T-DNA-RFD1* compound transcript with increased transcript size in *rfd1-1* initiating from the upstream T-DNA insertion (Fig. 1H, SFig. 1D) (Hedtke and Grimm, 2009). Notably, the endogenous *RFD1* mRNA isoform is not detected in *rfd1-1*. Our transcript analyses in *qua1-1* and *rfd1-1* are consistent with the hypothesis that initiation from the downstream gene promoter is repressed through the act of RNAPII transcription.

**Figure 1:**
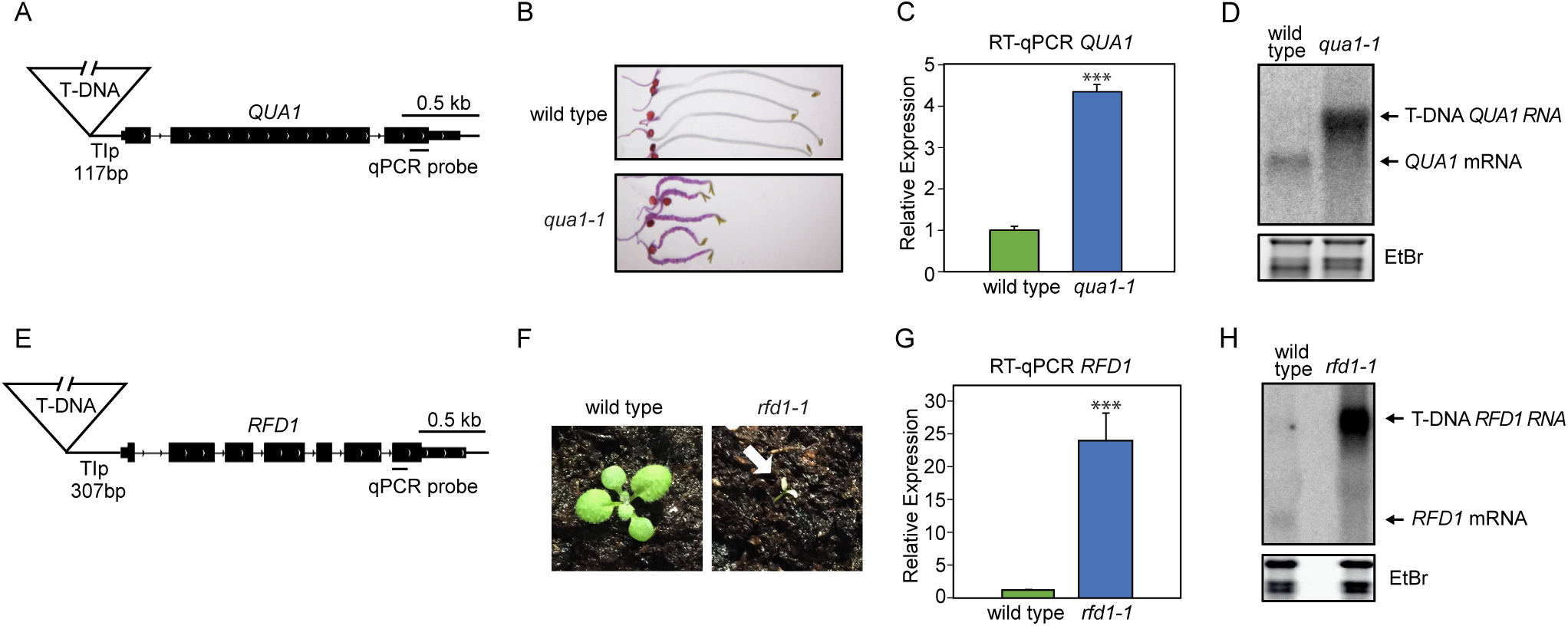
**Upstream transcription from within T-DNA represses downstream gene expression.** (A) Schematic representation of the *qua1-1* locus, including primer pair position for RT-qPCR. *Tip* denotes <u>T</u>ranscriptionally Interfered <u>p</u>romoter region remaining in the *qua1-1* mutant. (B) Ruthenium red staining of wild type and *qua1-1* hypocotyls (ecotype WS). (C) Quantitative analysis of *QUA1* transcript levels in wild type and *qua1-1* by RT-qPCR. Error bars represent standard deviation resulting from three independent replicates. Three asterisks denote p<0.001 between samples by Student’s t-test. (D) Analysis of *QUA1* transcript size and levels in wild type and *qua1-1* by northern blotting. Ethidium bromide (EtBr) staining of ribosomal RNA (rRNA) is used as a loading control. (E) Schematic representation of the *rfd1-1* locus, including primer pair position for RT-qPCR. (F) Photo-bleaching phenotype of *rfd1-1* seedlings grown in high light conditions (ecotype Col-0). (G) Quantitative analysis of *RFD1* transcript levels in wild type and *rfd1-1* by RT-qPCR. Error bars represent standard deviation resulting from three independent replicates. Three asterisks denote p<0.001 between samples by Student’s t-test. (H) Analysis of *RFD1* transcript size and levels in wild type and *rfd1-1* by northern blotting. EtBr staining of ribosomal RNA is used as a loading control.

To test if the genomic region between the *rfd1-1* T-DNA insertion and the translation start of *RFD1* can function as a promoter (designated as *TIp*_*RFD1*_, Fig. 1E; SFig. 1C) we assayed marker gene expression. We first monitored GUS-staining following transient expression in *Nicotiana benthamiana* and *Arabidopsis* leaves (Fig. 2A,B). We detect GUS activity in *TIp*_*RFD1*_-*GUS* compared to the p19 control injection in *N. benthamiana*, but it is weaker than the *p35S-GUS* positive control. We also detect stronger GUS staining and eYFP activity compared to p19 negative control injections in *Arabidopsis* (Fig. 2B, SFig. 2). To test if *TIp* can drive gene expression in relevant tissues and at sufficiently high levels, we performed a molecular complementation of the read-through mutants with genomic constructs driven by their respective *TIp*. We detect RFD1-FLAG protein expression to varying degrees in independent transformant lines by western blotting (Fig. 2C). Importantly, RFD1 expression driven by *TIp*_*RFD1*_-*RFD1-FLAG* complements the *rfd1-1* phenotype (Fig. 2D). Likewise, we detect QUA1-FLAG protein expression in independent *TIp*_*QUA1*_-*QUA1-FLAG* transformant lines by western blotting, and these lines complement the *qua1-1* phenotype (Fig. 2E,F). Thus, *TIp* DNA regions provide necessary and sufficient promoter activity to drive functional *RFD1* or *QUA1* expression. Interfering RNAPII transcription across *TIp* provides a candidate mechanism to explain the repression of initiation despite transcriptional activity at these regions.

**Figure 2:**
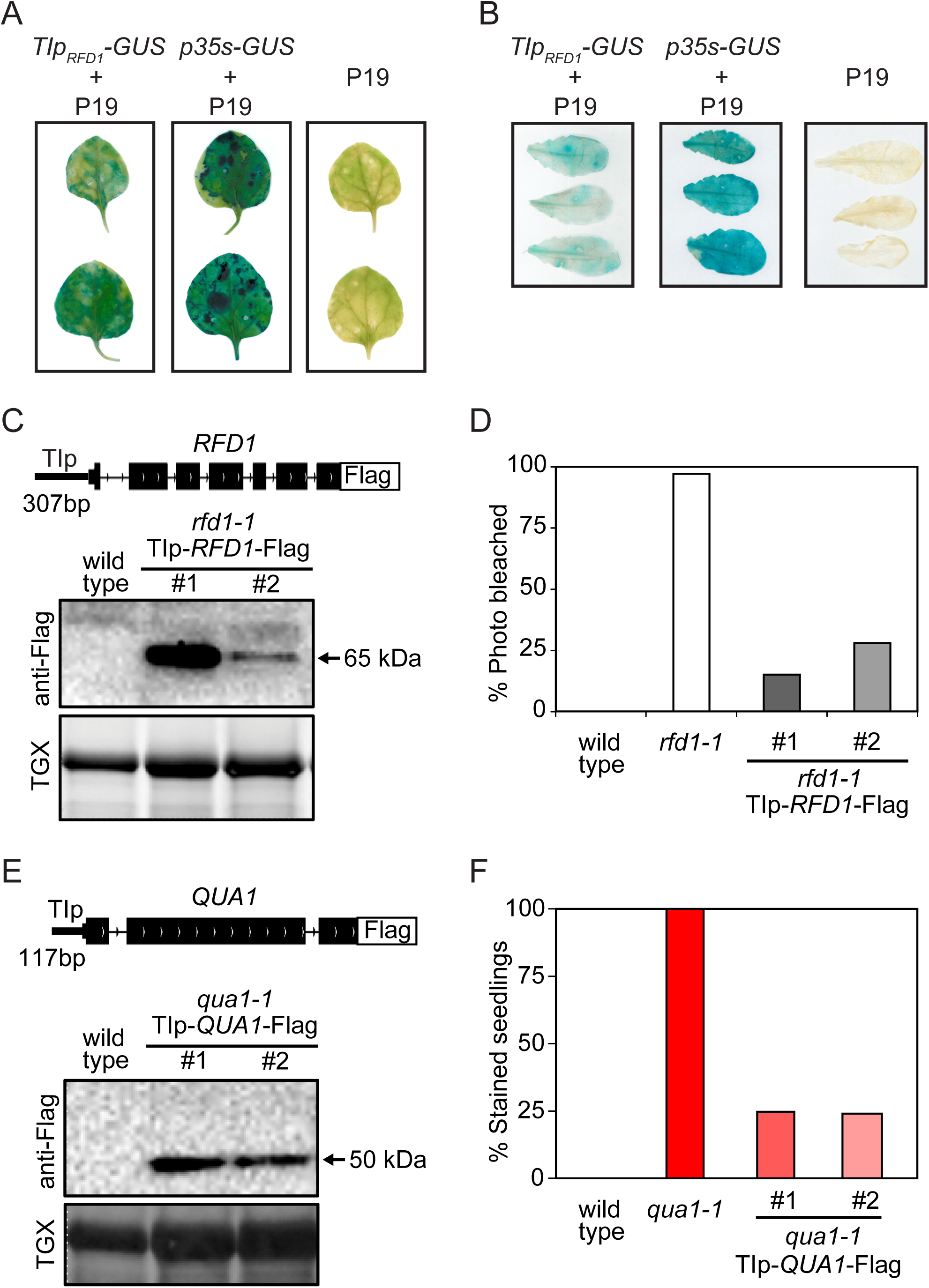
**The act of upstream transcription represses a functional downstream promoter.** (A) Transient transformation of GUS reporter gene under the control of *TIp*_*RFD1*_ in *N. benthamiana*. *p35s-GUS* and *p19* (lacking GUS) are positive and negative controls for GUS expression, respectively. (B) Transient transformation of GUS reporter gene under the control of *TIp*_*RFD1*_ in *Arabidopsis efr* mutant. *p35s-GUS* and *p19* (lacking GUS) are positive and negative controls for GUS expression, respectively. (C) Detection of RFD1-FLAG protein from *TIp*_*RFD1*_ by western blotting. For loading controls, total protein levels were detected using TGX stain-free protein gels. (D) Expression of RFD1-FLAG from *TIp*_*RFD1*_ complements *rfd1-1* photo-bleaching phenotype. Wild type (n=84), *rfd1-1* (n=142), #1 (n=1196), #2 (n=1142). (E) Detection of QUA1-FLAG protein from *TIp*_*QUA1*_ by western blotting. For loading controls, total protein levels were detected using TGX stain-free protein gels. (F) Expression of QUA1-FLAG from *TIp*_*QUA1*_ complements *qua1-1* phenotype. Wild type (n=97), *qua1-1* (n=96), #1 (n=267), #2 (n=254).

### Elevated RNAPII elongation signatures are found at promoters repressed through the act of upstream RNAPII transcription in Arabidopsis

Repressive RNAPII elongation across *TIp* in *qua1-1* and *rfd1-1* mutants may impact on molecular signatures associated with RNAPII elongation and initiation at *TIp*. To test this, we performed quantitative chromatin immunoprecipitation (qChIP) experiments for RNAPII initiation and elongation hallmarks. The elongating form of RNAPII (RNAPII-Ser2P) is enriched towards the 3’ end of the *QUA1* gene and depleted from the *QUA1* promoter in wild type *Arabidopsis* (SFig. 3A, B). H3K36me3 is enriched towards the 5’ end of genes in *Arabidopsis*, while H3K36me2 corresponds to the elongation phase and accumulates towards the 3’ end (Mahrez et al., 2016). We find the same pattern along the *QUA1* gene (SFig. 3C-D). Histone modifications of active promoters such as histone H3 acetylation (H3ac) and H3K4me3 are enriched towards the *QUA1* promoter (SFig. 3E-F) (Mahrez et al., 2016; Roudier et al., 2011; Zhang et al., 2016a). RNAPII initiation and elongation can be distinguished by our qChIP analysis.

**Figure 3:**
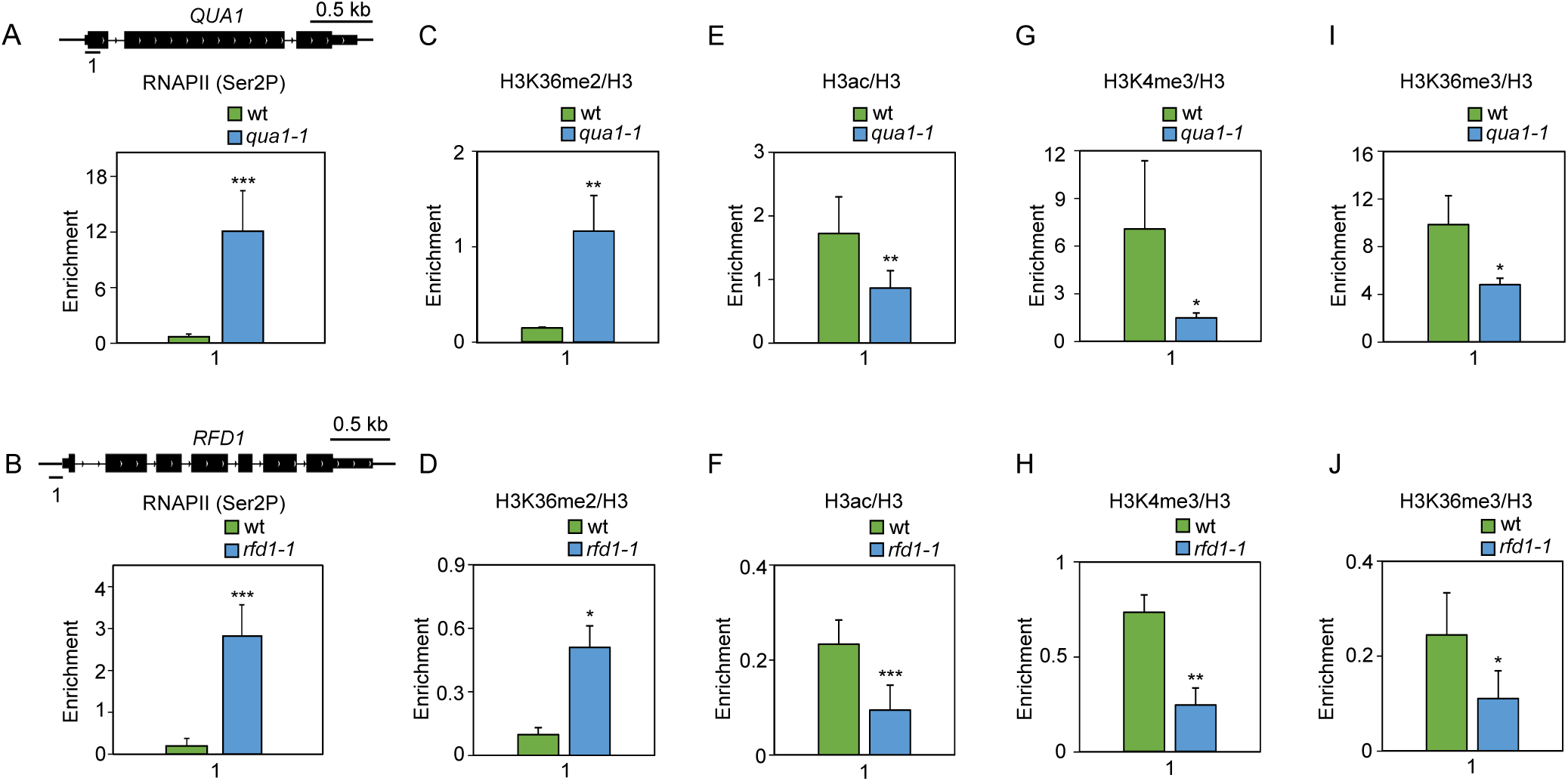
**Promoters repressed by upstream transcription adopt RNAPII elongation signatures.** (A, B) qChIP in mutants and their respective wild type ecotypes using promoter-proximal primer pairs for RNAPII (Ser2P). qChIP in mutants and their respective wild type ecotypes using promoter-proximal primer pairs for histone 3 (H3) modifications are shown (C-J). Data are normalized to bulk and show: (C, D) H3K36me2/H3, (E, F) H3 acetylation (H3ac/H3), (G, H) H3K4me3/H3, and (I, J) H3K36me3/H3. Error bars represent standard deviation resulting from at least three independent replicates. For statistical tests, a single asterisk denotes p<0.05, two asterisks denote p<0.01, three asterisks denote p<0.001 between samples by Student’s t-test.

We profiled *qua1-1* and *rfd1-1* mutants by qChIP to determine the impact of upstream RNAPII transcription across *TIp*_*QUA1*_ and *TIp*_*RFD1*_. Compared to their respective wild type ecotype, significantly higher levels of RNAPII-Ser2P were present at the position of promoter-proximal primer pairs in *qua1-1* and *rfd1-1* (Fig. 3A and 3B). These results support increased RNAPII elongation across the downstream promoter. Since bulk histone density remains largely unchanged across *QUA1* and *RFD1* in their respective mutants (SFig. 3G-I), we tested the presence of the elongation-specific chromatin signature H3K36me2. The mutants displayed increased levels of H3K36me2 at *TIp*_*QUA1*_ and *TIp*_*RFD1*_ (Fig. 3C and 3D). The increase of RNAPII elongation signatures at these promoters during repression indicates that these regions may now identify as zones of RNAPII elongation instead of promoters. Consistent with this hypothesis, histone modifications associated with active promoters (H3ac, H3K4me3, H3K36me3) were significantly depleted at *TIp*_*QUA1*_ and *TIp*_*RFD1*_ in the mutants (Fig. 3E-J). Collectively, these results demonstrate that upstream RNAPII transcription shifts the POINS to specify downstream promoters as intragenic regions. Our data suggest that promoter repression in these mutants is driven by transcription-mediated changes to promoter chromatin status.

### Arabidopsis FACT is required for gene repression through the act of upstream RNAPII transcription

Our analyses support that gene promoters can be repressed by interfering RNAPII elongation in *Arabidopsis*. We predict that factors associated with RNAPII elongation, such as the FACT complex, may be required for repression. To test the role of FACT in promoter repression by read-through transcription in *Arabidopsis*, we combined the previously described knock-down alleles of *spt16-1* and *ssrp1-2* mutants with *qua1-1* (Lolas et al., 2010). Ruthenium red staining comparing single and double mutants revealed patches of unstained hypocotyls in *spt16-1 qua1-1* compared to *qua1-1* (Fig. 4A). Importantly, *spt16-1 qua1-1* rescued the dwarf hypocotyl phenotype observed in *qua1-1* (Fig. 4B). These results indicate tightened cell-adhesion, and partial suppression of the *qua1-1* phenotype. The rescue effect was even more pronounced in *ssrp1-2 qua1-1* compared to *spt16-1 qua1-1* (Fig. 4A, B). Presumably this can be explained by a stronger knock-down of protein levels in *ssrp1-2* compared to *spt16-1* (Lolas et al., 2010). To test the link between H3K36me and FACT we assayed genetic interactions between *qua1-1* and a mutation in the *Arabidopsis* H3K36me methyltransferase *SDG8/ASHH2* (Grini et al., 2009; Zhao et al., 2005). Interestingly, we find no evidence for suppression of *qua1-1* by *sdg8-2* (SFig. 4). Together, our data demonstrate that the FACT complex is required for the cell adhesion defects observed in *qua1-1*.

**Figure 4:**
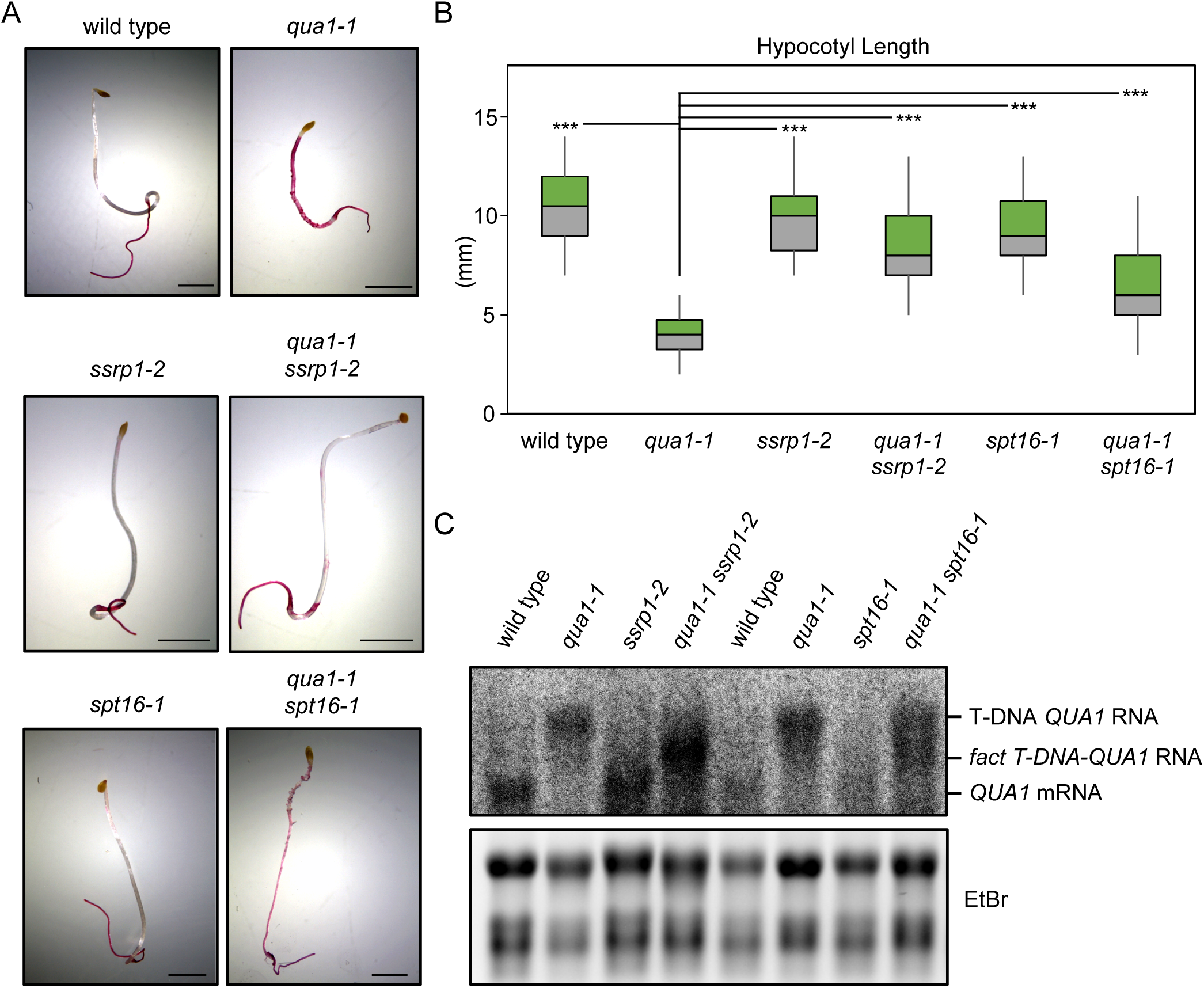
**The *Arabidopsis* FACT complex is required for downstream gene repression through the act of RNAPII transcription.** (A) Ruthenium red staining of wild type, *qua1-1, spt16-1, ssrp1-2*, and double mutants *qua1-1*/*spt16-1* and *qua1-1*/*ssrp1-2*. Scale bar represents 2 mm. (B) Quantification of hypocotyl length (mm) for 7 day, dark-grown wild type (n=30), *qua1-1* (n=30), *spt16-1* (n=30), *ssrp1-2* (n=30), and double mutants *qua1-1*/*spt16-1* (n=30) and *qua1-1*/*ssrp1-2* (n=30). Three asterisks denote p<0.001 between *qua1-1* and all other samples by Student’s t-test. (C) Analysis of *QUA1* transcripts in wild type, *qua1-1, ssrp1-2*, and the *qua1-1*/*ssrp1-2* double mutant as well as in *spt16-1* and the *qua1-1*/*spt16-1* double mutant by northern blotting. Ethidium bromide (EtBr) staining of ribosomal RNA (rRNA) is used as a loading control.

If phenotypic suppression of *qua1-1* through *fact* mutants was mechanistically linked to gene repression through the act of upstream interfering transcription, we would predict transcriptional changes. To examine the pattern of *QUA1* transcripts we performed northern blotting in single and double mutants. While the transcript pattern in *spt16-1* and *ssrp1-2* is indistinguishable from wild type controls, we observe new transcript patterns in *spt16-1 qua1-1* and *ssrp1-2 qua1-1* double mutants compared to *qua1-1* (Fig. 4C). Importantly, we detect a reduced size of the interfering transcript in *fact* mutants, arguing for a 5’-truncated transcript initiating from a cryptic TSSs. Notably, the wild type *QUA1* mRNA is not restored. This indicates that functional mRNAs are produced from an upstream promoter accessible in *fact* mutants. Our results support the conclusion that the activity of the FACT complex as part of RNAPII elongation suppresses intragenic TSSs.

### Arabidopsis FACT restricts the activity of intragenic promoters

To test if FACT suppresses endogenous intragenic TSSs, we measured *Arabidopsis* TSSs by 5’-CAP-sequencing (TSS-seq) (Pelechano et al., 2016). We called 77292 TSS clusters and annotated them by genomic location. Many TSS clusters (n=31013, or 40.1%) mapped to gene promoters (SFig. 5A). We obtained on average 47 million raw reads for two biological repeats of wild type, *spt16-1*, and *ssrp1-2* (Supplementary Table S1). Biological repeats showed a high degree of correlation (SFig. 5B). TSS-seq also revealed reduced TSS signals at *SPT16* and *SSRP1* genes in the respective mutants consistent with reduced expression (SFig. 5C-D). We examined the overlap of our TSS clusters with TSSs identified by CAGE (Cap Analysis Gene Expression) (Tokizawa et al., 2017). 74.8% of TSS clusters in core promoters overlap with at least one previously reported CAGE peak (SFig. 5E and Supplementary Table S2), indicating very good overlap across techniques and samples. Alternative mRNA isoforms of the *At4G08390* gene are differentially targeted to mitochondria or chloroplast, and our data resolves TSSs corresponding to these isoforms (SFig. 5F) (Obara et al., 2002). Interestingly, our TSS-seq data reveals almost as many TSSs in exons (n=30831, or 39.9%) as in gene promoters (Fig. 5A and Supplementary Table S3). In conclusion, these data illustrate high reproducibility of our TSS-seq methodology, and its abilities to validate TSSs as well as to reveal novel TSSs.

**Figure 5:**
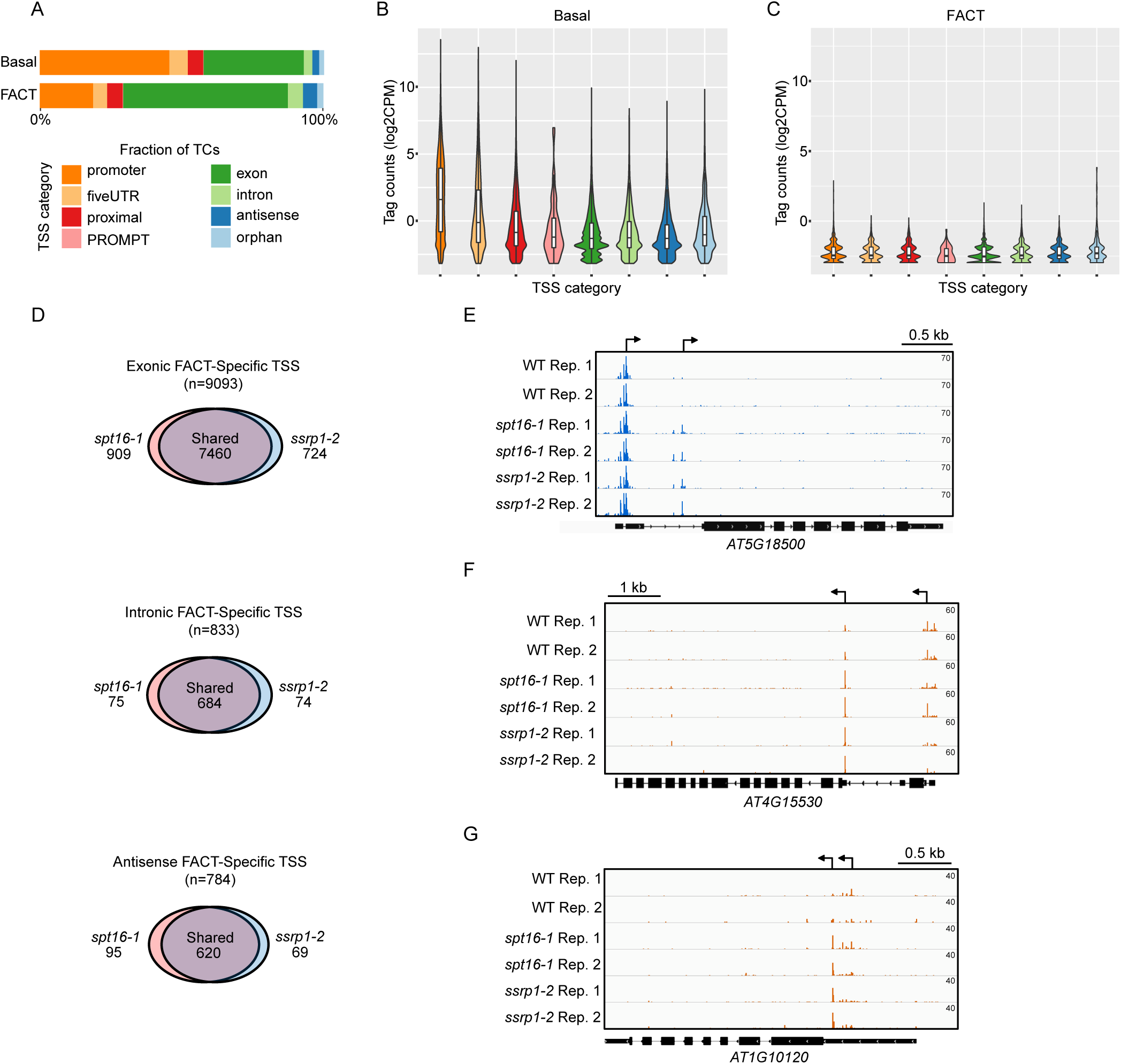
**FACT represses intragenic TSSs across the *Arabidopsis* genome.** (A) Genomic annotations of the basal set (upper) and the FACT-specific set of TSS (lower). (B, C) Distribution of log2-transformed expression values within each annotation category of the basal set (B) and the FACT-specific set of TSS (C). (D) Venn diagrams of exonic, intronic, antisense FACT-specific TSSs.(E) Screenshot of *fact* mutant-specific intragenic TSS observed in the *AT5G18500* gene. (F) Screenshot of alternative TSS for the *AT4G15530* gene that accumulates in *fact* mutants. (G) Screenshot of closely-spaced alternative TSSs observed for the *AT1G10120* gene.

To test the role of FACT in regulating TSSs in *Arabidopsis*, we divided the TSSs into three groups: i) constitutive TSSs detected in both wild type and FACT mutants (n=60966, or 78.9%); ii) wild-type specific TSSs (n=669, or 0.87%); and iii) TSSs specifically detected in *fact* mutants (n=15657, or 20.3%, Supplementary Table S4). The 23.4-fold increase of FACT-specific TSsS over the wild-type specific TSSs suggests that the FACT complex largely represses TSSs. FACT-specific TSSs map to intragenic locations, particularly exons (Fig. 5A). However, TSSs induced in *fact* mutants have a lower TSS-seq count compared to the basal TSS set indicating lower expression (Fig. 5B-C). The large majority of FACT-specific TSSs (7460 out of 9093, or 82%) were detected in both mutants (Fig. 5D). In addition, FACT-specific TSSs overlap less frequently with TSSs identified by CAGE (SFig. 5E and Supplementary Table S2). As much as 80.7% of FACT-specific exonic TSS clusters did not overlap with any CAGE peak. The *At5G18500* gene illustrates the induction of an intronic TSS in *fact* mutants (Fig. 5E). The *AT4G15530* pyruvate orthophosphate dikinase (PPDK) gene reveals preferential usage of a downstream TSS in *fact* mutants (Fig. 5F). While the longer transcript localizes to chloroplasts, the shorter PPDK isoform targeted to the cytoplasm instead (Parsley and Hibberd, 2006). The *AT1G10120* gene TSS selection demonstrates selective RNAPII initiation from a downstream 5’UTR site in *fact* mutants (Fig. 5G). The number of FACT-sensitive intragenic TSSs argues for high selectivity and tight regulation of intragenic regions functioning as TSSs. Our TSS-seq data therefore support a role for the FACT complex as part of POINS in *Arabidopsis*, with a specific function in suppressing intragenic cryptic TSSs.

### H3K4me1 in wild type marks intragenic regions that function as TSSs in FACT mutants

Common signatures in DNA sequence or chromatin environment may predispose intragenic regions to function as TSSs in *fact* mutants. We tested differential DNA-motif enrichment in exonic FACT-specific TSS clusters compared to basal exonic TSS. However, we detect no differentially enriched sequence motif or position bias (SFig. 5G). We next compared the chromatin signatures around exonic FACT-specific TSSs using available data for *Arabidopsis* in bud and leaf tissue samples (Zhang et al., 2016b). We compared chromatin signatures centered on five sets of genomic locations: FACT-specific exonic TSS, exonic TSSs not regulated by FACT (basal TSSs), promoter TSSs, random exonic positions and random genomic positions. Strikingly, we find the highest H3K4me1 enrichment at exonic sites in wild-type for FACT-specific TSSs (Fig. 6A-B). These regions also show less H3K27me3 compared to random exonic TSSs or random genomic locations (Fig. 6C-D). However, H3K27me3 is also reduced at basal exonic TSSs. If exonic TSS positions repressed by FACT were characterized by reduced chromatin condensation we would predict these sites to be particularly sensitive to DNaseI. Indeed, DNaseI sensitivity peaks upstream of promoter TSSs (Fig. 6E-F). However, we detect no specific DNaseI hyper-sensitivity compared to random genomic locations or basal exonic TSSs (Fig. 6E-F). Promoter TSSs are enriched for H3K27ac downstream of the TSSs (Fig. 6G-H). Low histone acetylation characterizes promoters repressed by overlapping RNAPII transcription (Fig. 3). While exonic regions carry higher H3K27ac than random exonic or random genomic positions, we detect no increase at FACT-specific exonic TSSs compared to basal exonic TSSs control regions (Fig. 6G-H). Our analyses of DNA elements and chromatin signatures link H3K4me1 to exonic regions poised to function as TSSs in *fact* mutants. Our results are consistent with a chromatin-based mechanism predisposing exonic sites as cryptic TSSs controlled by the FACT complex.

**Figure 6:**
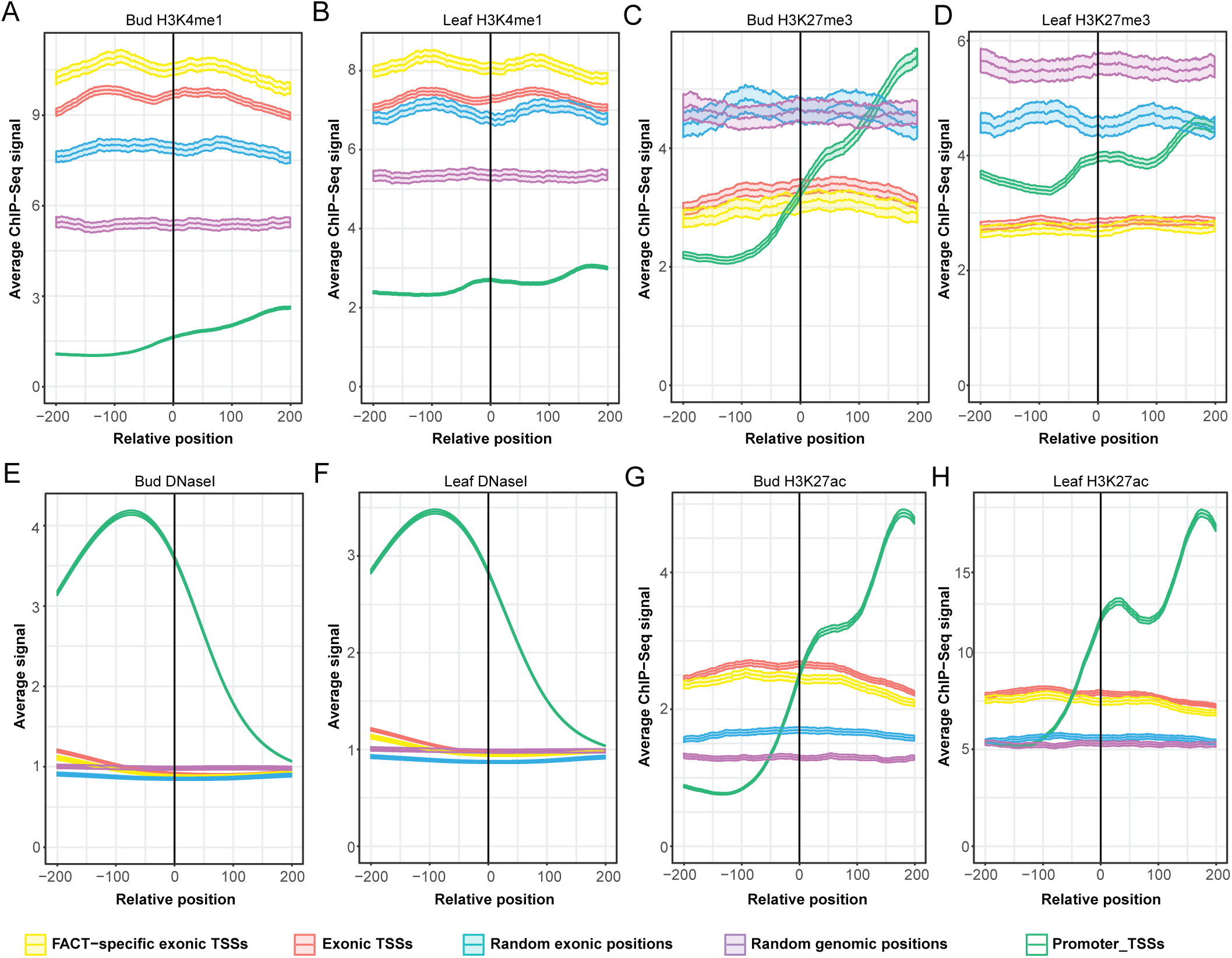
**H3K4me1 predisposes FACT-specific TSSs in exons.** Metagene profiles of chromatin signatures in 400 base pair intervals around FACT-specific exonic TSSs. A comparison to chromatin signatures derived from floral bud samples (A,C,E,G) and leaf samples (B,D,F,H) are shown. Data for H3K4me1 (A-B), H3K27me3 (C-D), DNaseI (E-F) and H3K27ac (G-H) are shown. (A-H) Data for FACT-specific exonic TSSs plotted in yellow (n=9043), data for the basal set of exonic TSSs plotted in red (n=21407), data for a random set of exonic positions plotted in blue (n=10000), data for a random set of genomic positions plotted in purple (n=10000) and data for promoter TSSs is plotted in green (n=31013). Data are shown with 95% confidence intervals.

All in all, our data support a model in which FACT is required for the repression of intragenic cryptic TSSs (Fig. 7). Read-through transcription can re-define gene promoters as intragenic to bring them under the influence of POINS. Repression of promoter function relies on a loss of initiation-specific RNAPII hallmarks, and a gain of elongation-specific signatures (Fig. 7A-B). The FACT complex represses intragenic transcription and maintains associated chromatin signatures (Fig. 7C). We identify a large number of exonic TSSs repressed throughout the *Arabidopsis* genome that are predisposed to function as TSSs by H3K4me1 in the absence of FACT-mediated repression. Our findings describe a conserved co-transcriptional chromatin-based mechanism shaping gene regulation and transcript isoform diversity.

**Figure 7:**
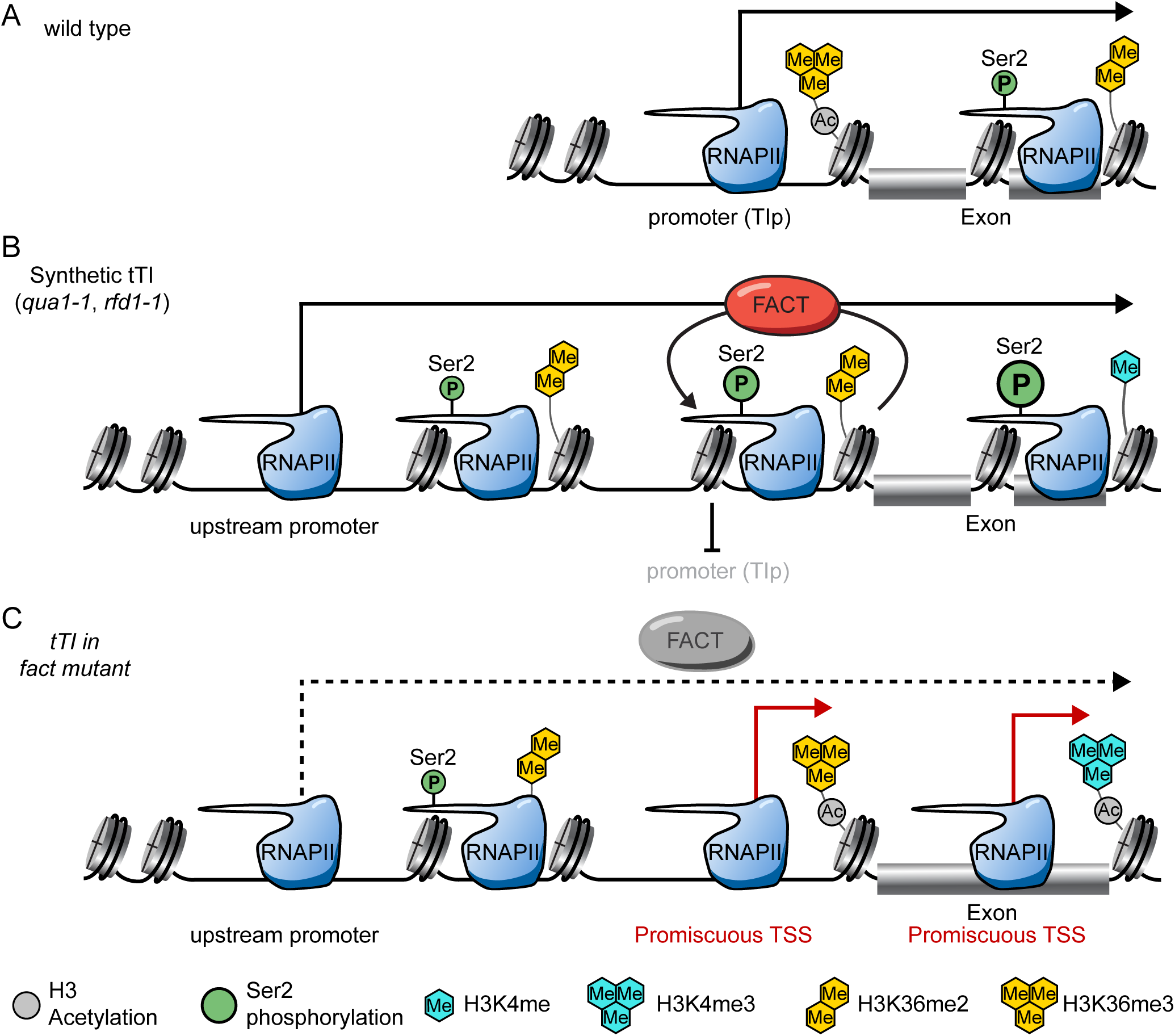
**Working model of how *fact* mutants suppress tandem transcription interference.** In wild type *Arabidopsis* RNAPII initiates transcription from promoter regions. Specific histone signatures such as H3 acetylation (grey circle) and H3K36me3 (yellow tri-hexagon) are associated with promoters, while RNAPII CTD Ser2 phosphorylation (green circle) and H3K36me2 (yellow di-hexagon) are associated with RNAPII elongation. (B) In *qua1-1* and *rfd1-1* mutants, upstream RNAPII transcription suppresses transcription initiation from downstream gene promoters (tandem Transcription Interference, tTI). The FACT complex contributes to tTI through chromatin remodeling activity. (C) Absence of FACT activity in *rfd1-1* and *qua1-1* mutants results in chromatin defects that trigger initiation of RNAPII transcription. H3K4me1 (turquoise) marks sites poised to function as intragenic TSSs in FACT mutants, presumably as H3K4me1 facilitates the transition to H3K4me3 that is linked to promoter activity. Intragenic RNAPII initiation in FACT mutants reduces tTI, as cryptic promoters may restore functional proteins.

## Discussion

TSSs shape RNA isoform expression but little is known about the mechanisms regulating TSS choice within transcription units. RNAPII transcription across gene promoters may re-define gene promoters as “intragenic” and repress them by mechanisms inhibiting initiation from within transcription units. We leveraged *Arabidopsis* T-DNA read-through mutants to identify a role of the conserved FACT histone chaperone complex in the repression of intragenic TSSs in a multicellular organism. Consistently, we identify a large number of intragenic TSSs controlled by FACT, particularly from exons with the chromatin signature H3K4me1.

Three activities of the FACT complex may explain a key role in repressing intragenic TSSs across species are: 1.) stimulation of RNAPII elongation, 2.) histone re-assembly in the wake of RNAPII transcription, and 3.) recycling of old histones to maintain POINS.

First, FACT stimulates RNAPII transcription of DNA templates packaged in nucleosome structures (Belotserkovskaya et al., 2003). Structural analyses suggest that the FACT complex directly binds nucleosomes on several contacts of histone proteins, stabilizing otherwise energetically unfavorable nucleosome conformations that weaken nucleosome binding to DNA (Hondele and Ladurner, 2013). Stabilization of partly unfolded nucleosome intermediates facilitates RNAPII progression through nucleosome barriers. Defective FACT may result in “transcription stress” through stalled or arrested RNAPII molecules in transcription units. Transcription stress serves as a DNA damage signal associated with proteolytic degradation of stalled RNAPII (Wilson et al., 2013). DNA damage-associated chromatin changes facilitate the initiation of RNAPII transcription that could help to explain elevated TSSs in FACT mutants (Francia et al., 2012; Price and D’Andrea, 2013). RNAPII elongation defects may hence trigger RNAPII initiation.

Second, FACT aids the re-assembly of nucleosomes in the wake of transcribing RNAPII (Belotserkovskaya et al., 2003; Schwabish and Struhl, 2004). Consistently, reduced nucleosome coverage within transcription units has previously been reported in human and yeast FACT mutants (Carvalho et al., 2013; van Bakel et al., 2013). Nucleosome Depleted Regions (NDRs) are associated with TSSs at gene promoters and may trigger firing from cryptic intragenic TSSs observed by us and others (Carvalho et al., 2013; Feng et al., 2016; Kaplan et al., 2003). However, we find no predisposition in wild type for relaxed chromatin packaging at exonic regions that function as TSSs in FACT mutants (Fig. 6). While it remains likely that NDRs are associated with intragenic TSSs in FACT mutants, these data suggest that NDRs would result from reduced FACT histone re-assembly activity rather than sequences of chromatin signatures destabilizing nucleosomes.

Third, the propensity of FACT to deposit “old histones” in the wake of RNAPII transcription represents an intuitive mechanism to maintain the co-transcriptional positional information provided by chromatin signatures (Jamai et al., 2009). Consistently, defective FACT disrupts POINS as is evidenced by the incorporation of the promoter-enriched histone variant H2A.Z within transcription units in FACT mutants (Jeronimo et al., 2015). Future studies will be required to dissect the individual and cumulative contributions of defects in RNAPII elongation, nucleosome re-positioning, and POINS establishment in the up-regulation of intragenic TSSs observed in FACT mutants.

Promoter-proximal histone acetylation is associated with active promoters. Histone deacetylases associate with elongating RNAPII to repress the activity of intragenic TSSs (Carrozza et al., 2005; Keogh et al., 2005). Our findings of reduced histone acetylation in promoter regions in *qua1-1* and *rfd1-1* read-through mutants support this observation in plants (Fig. 3). We find no evidence for a predisposition to increased H3K27ac at exonic regions in wild type that function as FACT-specific TSSs (Fig. 6). These data argue that reduced histone acetylation within gene-bodies results from the process of RNAPII elongation, although different specificities of histone acetylation antibodies could also contribute to this observation. Proteomic screens identified several histone deacetylases (HDACs) associating RNAPII elongation complexes in *Arabidopsis* (Antosz et al., 2017). It remains to be tested which complexes provide HDAC activity linked to the suppression of intragenic TSSs. H3K36me3 correlates with RNAPII elongation and linked suppression of intragenic TSSs in mammals and yeast. However, H3K36me3 is a promoter-proximal chromatin signature in *Arabidopsis*, while H3K36me2 co-localizes with RNAPII elongation (Mahrez et al., 2016). Consistently, our chromatin state analyses in *qua1-1* and *rfd1* support H3K36me2 as a chromatin signature of RNAPII elongation, and H3K36me3 as a signature associated with initiation zones near promoters. Interestingly, we find no evidence for a role of the *Arabidopsis* H3K36 methyltransferase *SDG8/ASHH2* is gene repression through the act of RNAPII transcription in *qua1-1* (SFig. 4). Perhaps one of the other 47 SET-Domain Genes (SDGs) in *Arabidopsis* contributes a redundant activity to the repress of intragenic TSSs (Ng et al., 2007). Future research will address the question if H3K36me2 fulfils the role of H3K36me3 in plants, or if mechanisms repressing intragenic TSSs independent of H3K36me exist. Our screen for chromatin signatures characterizing exonic regions poised to function as TSSs in FACT mutants identifies histone 3 lysine 4 mono-methylation (H3K4me1). The association of H3K4me1 with enhancer activity has recently been called into question (Dorighi et al., 2017), however H3K4me1 is indicative of RNAPII transcription. In *Arabidopsis*, intragenic H3K4me1 counteracts H3K9me2-mediated chromatin repression (Inagaki et al., 2017). High levels of H3K4me1 at FACT-respressed TSS positions may facilitate the transition towards H3K4me3 and promoter function of exonic regions. Further dissection of the regulation of intragenic promoters in plants promises to reveal novel mechanisms for the co-transcriptional HDAC recruitment, involved histone methyltransferases and novel components of chromatin-based suppression of intragenic TSSs.

Alternative TSS selection provides a mechanism to generate transcript isoform diversity from a given genome. Light signaling triggers the choice between alternative TSSs in *Arabidopsis*, with profound effects on subcellular protein targeting (Ushijima et al., 2017). While a role of FACT in plant light signaling remains to be firmly established, it is conceivable that FACT contributes to the regulation of light-sensitive intragenic TSS selection. TSSs at gene promoters identified by our TSS-seq approach agrees well with promoter TSSs identified by CAGE, despite the fact that CAGE data used for comparison was sampled using different developmental stages (Tokizawa et al., 2017). However, our TSS-seq data in FACT mutants identify thousands of novel TSSs mapping to intragenic regions, highlighting that FACT activity likely protects plant genomes from intragenic transcription initiation. While our bioinformatic methods to detect intragenic TSSs are rigorous, and we identify a large overlap between TSSs in mutants of two FACT subunits, it is clear that expression of FACT-specific TSSs is low compared to basal TSSs (Fig. 5). Since null mutations in FACT are not viable, we characterized knock-down alleles. This suggests we are underestimating the global impact of FACT activity on intragenic TSS suppression, both in in quality and quantity (SFig. 5) (Lolas et al., 2010). Moreover, an low expression of *fact*-specific TSSs may be explained by targeted RNA degradation, as has been shown for cryptic expression of divergent lncRNA transcripts (Ntini et al., 2013). *Arabidopsis spt16-1* and *ssrp1-2* display similar phenotypic defects, indicating that regulation of intragenic TSSs may shape plant development. Moreover, an increasing number of examples of gene regulation by acts of interfering lncRNA transcription requiring FACT are emerging (Ard et al., 2017). While specific examples remain to be characterized in plants, we demonstrate that the underlying mechanism of repressive RNAPII transcription is operational.

Future research will be required to fully address potential regulatory functions of intragenic TSSs. Our study offers a platform to query the role of intragenic TSSs in plant signalling and development, both by providing conceptual advances for a mechanism regulating intragenic TSSs, and by the identification of thousands of exonic sites poised to function as TSSs by H3K4me1. Our characterization of conserved players in repressive RNAPII transcription will inform efforts to characterize gene regulation through the act of pervasive lncRNA transcription in plants and beyond.

## Acknowledgments

We thank Jasmin Dilgen for technical assistance, Grégory Mouille and Henning Mühlenbeck for help with the ruthenium red staining assay, Jan Høstrup for plant care, Laura Brey for help in initial stages of the project, Albin Sandelin, Malte Thodberg and Axel Thieffry for help with TSS data analysis, and members of the S.M. laboratory for critical reading of the manuscript. Research in the laboratory of S.M. is supported by a Hallas-Møller Investigator award by the Novo Nordisk Foundation, a Copenhagen Plant Science Centre Young Investigator Starting grant and the ERC StG2017-757411.R.A. is supported by an EMBO LTF (ALTF 463-2016). P.K. is supported by MSCA-IF 703085. V.P. is supported by a SciLifeLab Fellowship, the Swedish Research Council (VR 2016-01842), a Wallenberg Academy Fellowship (KAW 2016.0123) and the Ragnar Söderberg Foundation.

The authors declare that they have no conflict of interest.

## Methods

### Plant Growth

All *Arabidopsis thaliana* lines used in this study are in the Col-0 ecotype background, with the exception of the *qua1-1* mutant, which is in WS ecotype background. Plants were grown in greenhouses or climate chambers with a 16h light/8h dark cycle at 22°C for general growth and seed harvesting. For seedlings grown on plates, the sterilized seeds were grown on 1/2 Murashige and Skoog (MS) medium containing 1% sucrose and supplemented with 0.5% Microagar. Information about T-DNA insertion mutant *Arabidopsis* lines is provided in Key Resource Table.

### Growing *rfd1-1* plants

For analysis of the homozygous *rfd1-1* phenotype, seeds were sown in 96 well trays stratified for 2-3 days at 4°C. F2 analysis trays were grown in high light conditions (> 100 μE). White seedlings were counted 10 days later. To propagate *rfd1-1* homozygotes, heterozygous *rfd1-1* seeds were sterilized and sown on MS plates with phosphinotricin selection, covered in foil, and stratified for 2 days at 4°C. Seeds were light induced for 6-8 h in a growth chamber with light strength of 80-100 μE. Plates were covered in foil for 3 days, the plates were unwrapped and grown in low light (< 50 μE) for 3-4 weeks before transferring to soil. To isolate RNA, *rfd1-1* homozygote seeds and corresponding wild type controls were sterilized and sown on MS plates as described above and grown in low light for two weeks. In order to collect enough material for ChIP, heterozygous *rfd1-1* seeds were sterilized and sown on MS plates with phosphinotricin selection as described above and grown in low light for two weeks. Col-0 wild type controls were treated the same way, but without selection, for corresponding.

### Ruthenium red staining

Seeds were sown in a 96 well plates containing 70 ul water. To synchronize germination, seeds were stratified at 4°C for 2-3 days. Germination of seeds was induced by light for 8-10 hours. The plates were wrapped in aluminium foil for 7 days. Etiolated seedlings were stained with 0.05% ruthenium red solution for 2 minutes. Seedlings were washed twice with water. Staining phenotype was recorded using a stereomicroscope.

### Cloning and plant transformation

Marker gene constructs were generated using pGWB vectors (Nakagawa et al., 2007). TIp*_RFD1_* was amplified from *rfd1-1* genomic DNA using primers MLO414/422. p35S was amplified from *rfd1-1* genomic DNA using primers MLO538/MLO416. TIp*_RFD1_* and p35S were inserted into pENTR-D-Topo vector through topo reaction to generate entry vector SMC358 (containing TIp*_RFD1_*) and SMC379 (containing p35S). Entry vectors were used in a LR reaction with pGWB533 (containing GUS) and pGWB540 (containing eYFP) to generate expression vector SMC371 (TIp*_RFD1_*-GUS), SMC367 (TIp*_RFD1_*-eYFP), SMC377 (p35S-GUS) and SMC373 (p35S-eYFP). The expression vectors were transformed into *Agrobacterium tumefaciens* strain GV3850 by electroporation under 2.5kV, 400Ω resistance and 25uF capacitance. *Agrobacteria* harboring expression vectors were respectively co-infiltrated with the p19 suppressor of silencing into *Nicotiana benthamiana* and *Arabidopsis thaliana efr* mutant leaves (Zipfel et al., 2006). GUS and eYFP signal was detected at 2 days after infiltration. Complementation constructs were generated using SMC330, a version of pEG302 (Earley et al., 2006) enabling Hygromycin selection following plant transformation. SMC330 was generated by replacing the phosphinotricin herbicide resistance gene with the Hygromycin resistance gene of pCambia1300. *TIp*_*QUA1*_:*QUA1* and TIp*_RFD1_:RFD1* were amplified from genomic wild type DNA using primers MLO727/728 and MLO414/442, respectively. The resulting PCR products were introduced into pENTR-D-Topo by topo cloning to generate entry vectors (SMC409 for *TIp*_*QUA1*_:*QUA1* and SMC356 for TIp*_RFD1_:RFD1*). The entry vectors were used in a LR reaction with SMC330 to generate expression vector SMC410 (containing *TIp*_*QUA1*_:*QUA1-FLAG* construct) and SMC380 (containing TIp*_RFD1_*:*RFD1-FLAG* construct). The complementation constructs were then transformed into *Agrobacterium tumefaciens* strain GV3101 (pMP90) by electroporation under 2.5kV, 400Ω resistance and 25μF capacitance. The complementation assay was done using *Agrobacterium*-mediated transformation described in (Clough and Bent, 1998). Homozygous *qua1-1* and heterozygous *rfd1-1 Arabidopsis* were used for complementation. Seeds from transformed *Arabidopsis* were screened for T-DNA integration by hygromycin resistance. Multiple independent single-locus insertions were identified by segregation analysis and tested for complementation and protein expression (Fig. 2).

### GUS Staining and Fluorescence Imaging

The GUS staining assay was performed as previously described (Jefferson et al., 1987). X-Gluc substrate was vacuum infiltrated into *Arabidopsis* and *N. benthamiana* leaves. After staining, leaves were rinsed in 70% ethanol at room temperature until the chlorophyll was washed off. YFP fluorescence was quantified using BIO-RAD imager Gel Doc.

### Western blotting

Equal amounts of plant material were harvested from plant tissue. Proteins were extracted in 2.5x extraction buffer (150 mM *Tris-HCl pH 6.8; 5 % SDS; 25 % Glycerol; 0.025 % Bromphenolblue; 0.1 mM DTT)*. Proteins were separated by SDS-PAGE using precast 8-16 % Criterion TGX stain-free protein gels (Biorad) and transferred to PVDF membrane using a semi-dry Trans-blot Turbo transfer system (Biorad). Membranes were blocked (5% non-fat dried milk in PBS) for 1 hour at RT. anti-FLAG (Sigma F3165) was added overnight at 4°C with rotation. Membranes were washed with PBS before the addition of the anti-mouse HRP-conjugated secondary antibody (Dako P0161) for 1 hour at room temperature. Membranes were washed in PBST. Chemiluminescent signals were detected using Super-Signal West Pico Chemiluminescent (ThermoFisher) according to manufacturer’s instructions.

### Quantitative chromatin immunoprecipitation

qChIP experiments were performed essentially as described (Marquardt et al., 2014), with minor modifications. For immunoprecipitations, Protein A magnetic beads (GenScript) and 2 ug of an antibody (Anti-Histone H3, ab1791; Anti-RNA polymerase II CTD YSPTSPS phosphor S2, ab5095; Anti-Histone H3 tri methyl K4, ab8580; Anti-Histone H3 tri methyl K36, ab9050; Anti-Histone H3 di methyl K36, a9049; Anti-Histone H3 pan-acetyl, ab47915) were added to solubilized chromatin. Quantitative analysis was performed on captured DNA by qPCR (Biorad). See Supplementary Tables S5 for oligonucleotide sequences. ChIP enrichments were calculated as the ratio of product of interest from IP sample to the corresponding input sample. For *qua1-1* and corresponding wild type (ecotype WS), results were further normalized to an internal reference gene (*ACT2*). Error bars represent standard deviation resulting from at least three independent replicates.

### RT-qPCR

RNA was isolated from 14 day old seedlings using Plant RNeasy Mini-Kits as per manufacturer’s instructions (Qiagen). For RT„qPCR experiments, first strand complementary DNA synthesis was performed on Turbo DNase-treated (Ambion) RNA using oligo-dT primers and Superscript III (Invitrogen) as per manufacturer’s instructions. Negative controls lacking the reverse transcriptase enzyme (-RT) were performed alongside all RT„qPCR experiments. Quantitative analysis was performed by qPCR (Biorad). See Supplementary Table S5 for oligonucleotide sequences. Data was normalized to an internal reference gene (*ACT2*). Levels in mutants represent relative expression compared to corresponding wild type.

### Northern blotting

Northern analyses were essentially performed as previously described with only minor modifications (Marquardt et al., 2011). Briefly, 5 micrograms of total RNA were separated by electrophoresis on agarose-formaldehyde-MOPS gels and transferred to a nylon transfer membrane by capillary blotting in 10x SSC overnight. RNA was crosslinked to the nylon membrane by UV irradiation. Membranes were probed with single stranded cDNA probes generated by incorporation of radioactive α-32P-dTTP. A Typhoon phosphoimager (GE Healthcare Life Sciences) was used for analysis.

### TSS-seq Library Construction

TSSs were mapped genome-wide in *Arabidopsis* using 5’-CAP-sequencing as previously described (Pelechano et al., 2016). Briefly, 5 micrograms of DNase-treated total RNA were treated with CIP (NEB) to remove all non-capped species in the sample. Next, 5’ caps were removed using Cap-Clip (CellScript) to permit ligation of single-stranded rP5_RND adapter to 5’-ends of previously capped species with T4 RNA ligase 1 (NEB). Poly(A)-enriched ligated RNAs were captured with oligo(dT) Dynabeads (Thermo Fisher Scientific) according to manufacturer’s instructions and fragmented in fragmentation buffer (50 mM Tris acetate pH 8.1, 100 mM KOAc, 30 mM MgOA) for 5 mins at 80°C. First-strand cDNA was generated using SuperScript III (Invitrogen) and random primers following manufacturer’s instructions. Second-strand cDNA was generated using Phusion high-fidelity polymerase (NEB) and the BioNotI-P5-PET oligo as per manufacturer’s instructions. Biotinylated PCR products were captured by streptavidin-coupled Dynabeads (Thermo Fisher Scientific), end repaired with End Repair Enzyme mix (NEB), A-tailed with Klenow fragment exo-(NEB), and ligated to barcoded Illumina compatible adapter using T4 DNA ligase (NEB). Libraries were amplified by PCR, size selected using AMPure XP beads (Beckman Coulter), pooled following quantification by bioanalyzer, and sequenced in single end mode on the following flowcell: NextSeq^®^ 500/550 High Output Kit v2 (75 cycles) (Illumina).

### Bioinformatic analysis

Quality of raw NGS data was consistently high as reported by the FastQC software (<https://www.bioinformatics.babraham.ac.uk/projects/fastqc/).> FASTQ files were subjected to quality and adapter trimming at 3’ ends using Trim Galore v0.4.3 (--adapter ATCTCGTATGCCG) (https://github.com/FelixKrueger/TrimGalore). UMI barcodes (8 nt) were trimmed from 5’ ends and appended to FASTQ headers using a custom Python script. The adapter-and UMI-trimmed reads were aligned to TAIR10 genome assembly using STAR v2.5.2b (--outSAMmultNmax 1 -- alignEndsType Extend5pOfRead1) (Dobin et al., 2013). The output SAM files were sorted and converted to BAM using Samtools v1.3.1 (Li et al., 2009). Reads aligned to rRNA, tRNA, snRNA or snoRNA loci were filtered out using BEDTools v2.17.0 (Quinlan and Hall, 2010). The resultant BAM files were filtered for reads with MAPQ ≥ 10 using Samtools. Finally, BAM files were deduplicated using a custom Python script (namely, groups of reads with 5’ ends aligned to the same genomic position and having identical UMI sequences were collapsed to a single representative read). The clean BAM files were converted to stranded Bedgraph files using BEDTools genomecov (-bg −5 - strand + for forward strand, -bg −5 strand - for reverse strand). Bedgraph files were compressed to BigWig format using kentUtils bedGraphToBigWig (https://github.com/ENCODE-DCC/kentUtils). The rest of the data analysis was done in R using Bioconductor packages. Transcription start sites (TSS) were called from BigWig files using the new CAGEfightR pipeline (https://github.com/MalteThodberg/CAGEfightR). Only genomic positions supported by at least two 5’ tags in at least two libraries from the same genotype were considered as TSS candidates. Adjacent TSS separated by not more than 20 bp were merged together into TSS clusters. The TSS clusters were annotated by intersection with various genomic features which were extracted from the TxDb.Athaliana.BioMart.plantsmart28 package. The package contains annotations from ENSEMBL Plant version 28 which combines TAIR10 and Araport11. In particular, proximal upstream regions were defined as [TSS-500bp, TSS-100bp], promoters as [TSS-100bp, TSS+100bp] and PROMPTs (promoter upstream transcripts) as intervals antisense to promoters. TSSs were annotated by genomic location as either genic (promoter, proximal, PROMPT, fiveUTR), intragenic (exon, intron, antisense), or intergenic (“orphan”). In case of conflicting annotations, a single annotation was chosen according to the following hierarchy: orphan < antisense < intron < exon < fiveUTR < PROMPT < proximal < promoter. Intersections between genomic coordinates were analyzed using methods from the GenomicRanges package (Lawrence et al., 2013) Metagene profiles were plotted using custom R scripts and the ggplot2 library. ChIP-Seq data for metagene analysis of histone modifications and DHS sites were downloaded as BigWig files from the PlantDHS database (Zhang et al., 2016b). Genomic DNA sequences around TSS summits were extracted using the BSgenome.Athaliana.TAIR.TAIR9 package. The differential motif enrichment analysis was done using the DREME software.

---

**Supplementary Figure 1:**
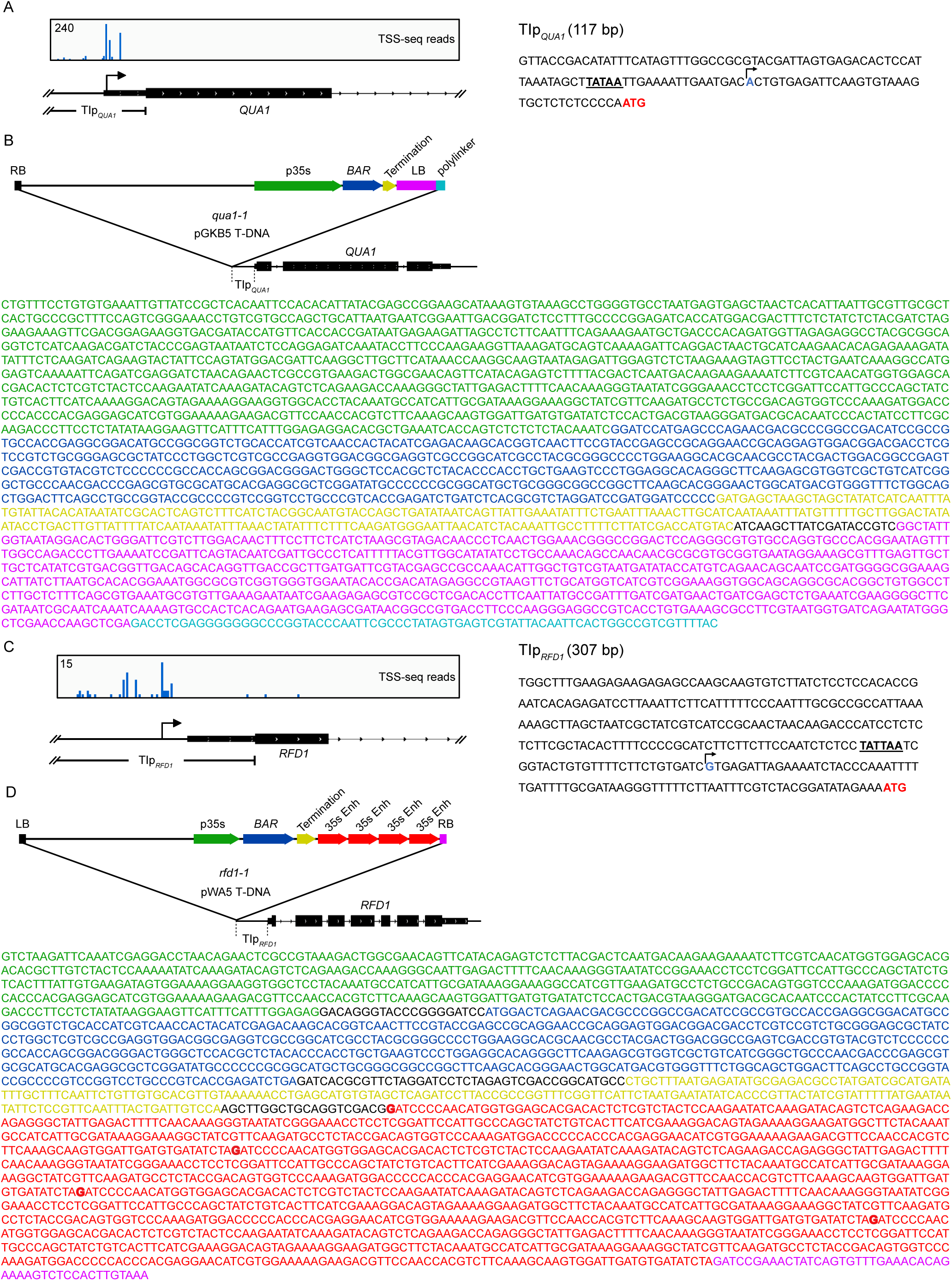
***RFD1* and *QUA1* promoters.** (A) The 117 bp *TIp*_*QUA1*_ promoter in *qua1-1* contains the *QUA1* TSS (as detected by TSS-seq in wild type) and upstream TATA element (bold and underlined). The predominant TSS peak is highlighted in blue. The start codon is highlighted in red.Detailed annotation of functional elements from p35s in *qua1-1* T-DNA insertion. Schematic diagram is given. BAR (Bialaphos Resistance) annotates the ORF conferring resistance to the plant herbicide Phosphinotricin. (C) The 307 bp *TIp*_*RFD1*_ promoter in *rfd1-1* contains the *RFD1* TSS (as detected by TSS-seq in wild type) and upstream TATA-like element (bold and underlined). The predominant TSS peak is highlighted in blue. The start codon is highlighted in red. (D) Detailed annotation of functional elements from p35s in *rfd1-1* T-DNA insertion. Schematic diagram is given, corresponding DNA sequence derived from sanger sequencing of genomic DNA in matching color is given below. BAR (Bialaphos Resistance) annotates the ORF conferring resistance to the plant herbicide Phosphinotricin. A tetrameric repeat of the 35S enhancer (35S Enh) sequence is located near the T-DNA right border (RB).

**Supplementary Figure 2:**
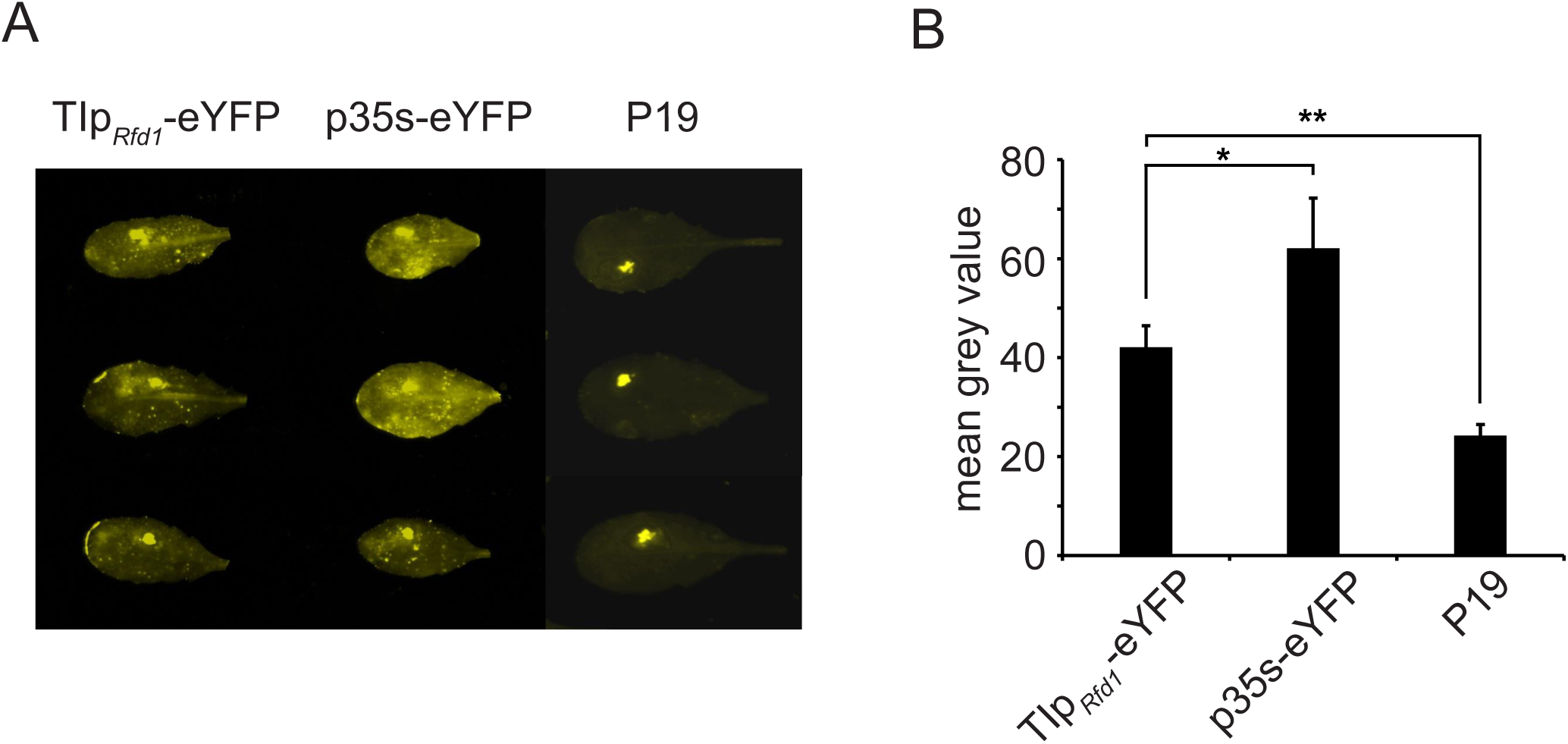
***TIp*_*RFD1*_ drives eYFP reporter gene expression in *Arabidopsis***. (A) Transient transformation of *eYFP* reporter gene under the control of *TIp*_*RFD1*_ in *Arabidopsis efr* mutant. *p35s-eYFP* and *p19* (lacking *eYFP*) are positive and negative controls for eYFP expression, respectively. (B) Quantification of eYFP signal in panel A using ImageJ based on three replicates of three infiltrated leaves per construct. * denotes p<0.05 and ** denotes p<0.01 between samples by Student’s t-test.

**Supplementary Figure 3:**
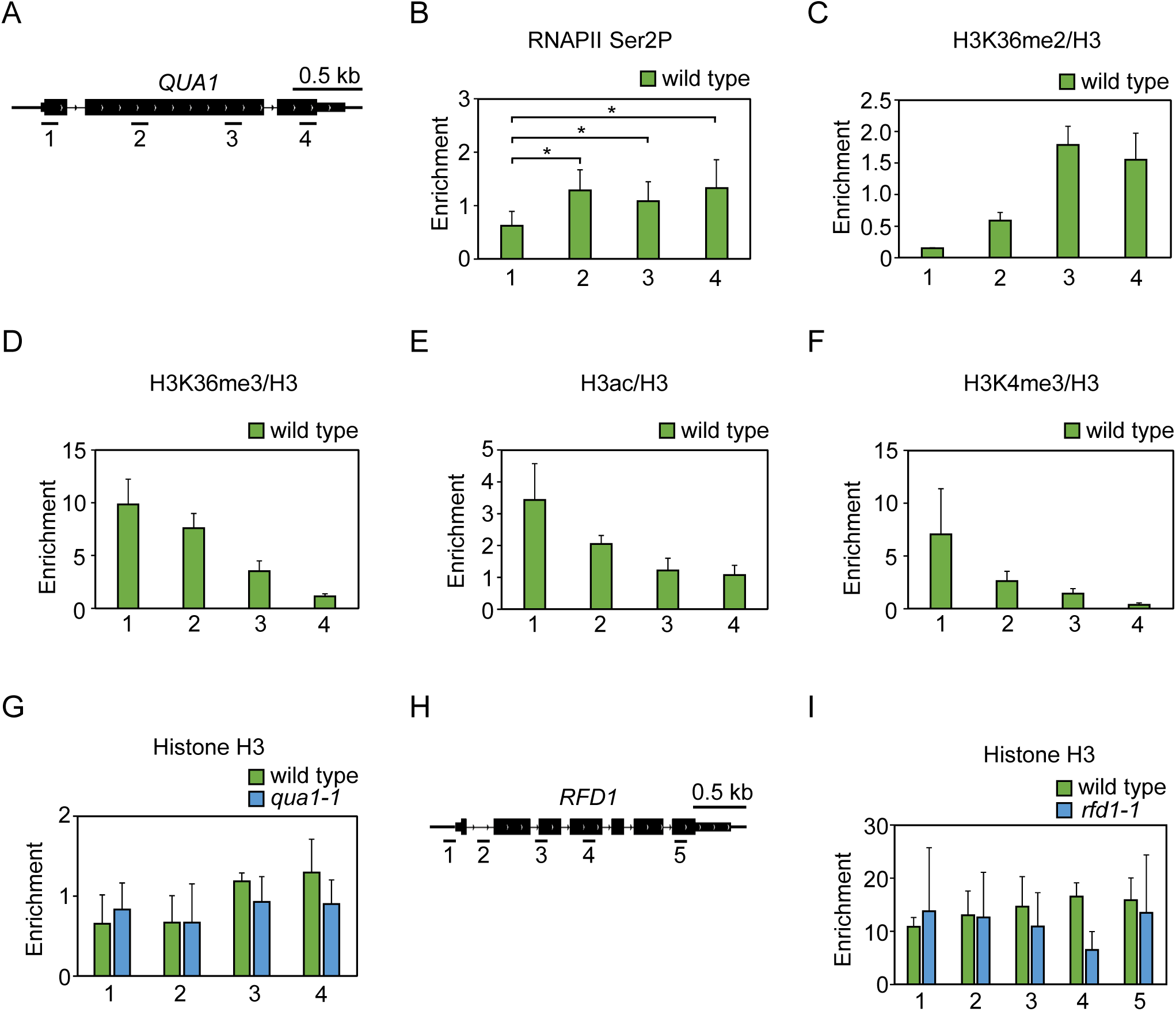
**Technical controls for qChIP analyses.** (A) Schematic representation of the *qua1-1* locus, including position of primer pairs for qChIP across *QUA1* gene in wild type (ecotype WS). (B) RNAPII (Ser2P) data across *QUA1* gene in wild type. For statistical tests, a single asterisk denotes p<0.05 between samples by Student’s t-test. qChIP across *QUA1* gene in wild type for (C) H3K36me2/H3, (D) H3K36me3/H3, (E) H3ac/H3 and (F) H3K4me3/H3. (G) Histone H3 qChIP across *QUA1* gene in wild type (ecotype WS) and *qua1-1*. (H) Schematic representation of the *rfd1-1* locus, including position of primer pairs. (I) Histone H3 qChIP across *RFD1* gene in wild type (ecotype Col-0) and *rfd1-1*. Error bars represent standard deviation resulting from at least three independent replicates.

**Supplementary Figure 4:**
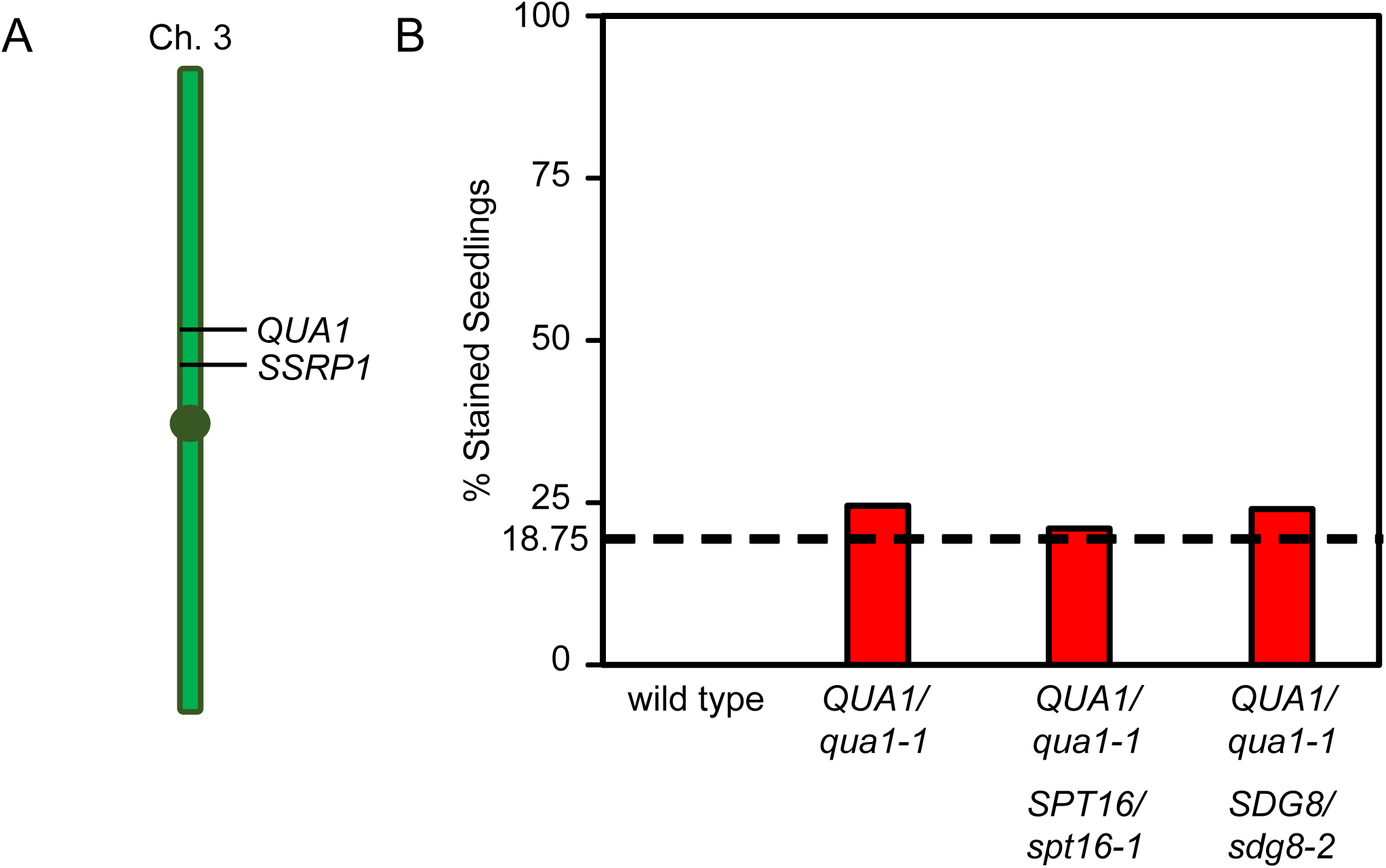
***qua1-1* suppression by *spt16-1*, but not *sdg8-2***. (A) The *QUA1* and *SSRP1* loci are linked on *Arabidopsis* chromosome 3. Genetic linkage prevents an analysis of *qua1-1* suppression by *ssrp1-2* using segregating populations. (B) Segregation analysis of *qua1-1*. We expect ¼ of the progeny from a *qua1-1/QUA1* parent to be *qua1-1/qua1-1* and hence stain by ruthenium red. Independently segregating loci that suppress *qua1-1* are expected to reduce the number of *qua1-1* mutants by a quarter (1/4 „ 1/16 = 18.75%) indicated by a dashed line. Based on the expected pattern of phenotypic segregation the FACT mutant *spt16-1* suppresses the *qua1-1* phenotype, while the H3K36 methyltransferase mutant *sdg8-2* does not. Wild type (n=97), *qua1-1/QUA1* (n=456), *qua1-1/QUA1*; *SPT16/spt16-1* (n=349), and *qua1-1/QUA1*; *SDG8/sdg8-2* (n=479).

**Supplementary Figure 5:**
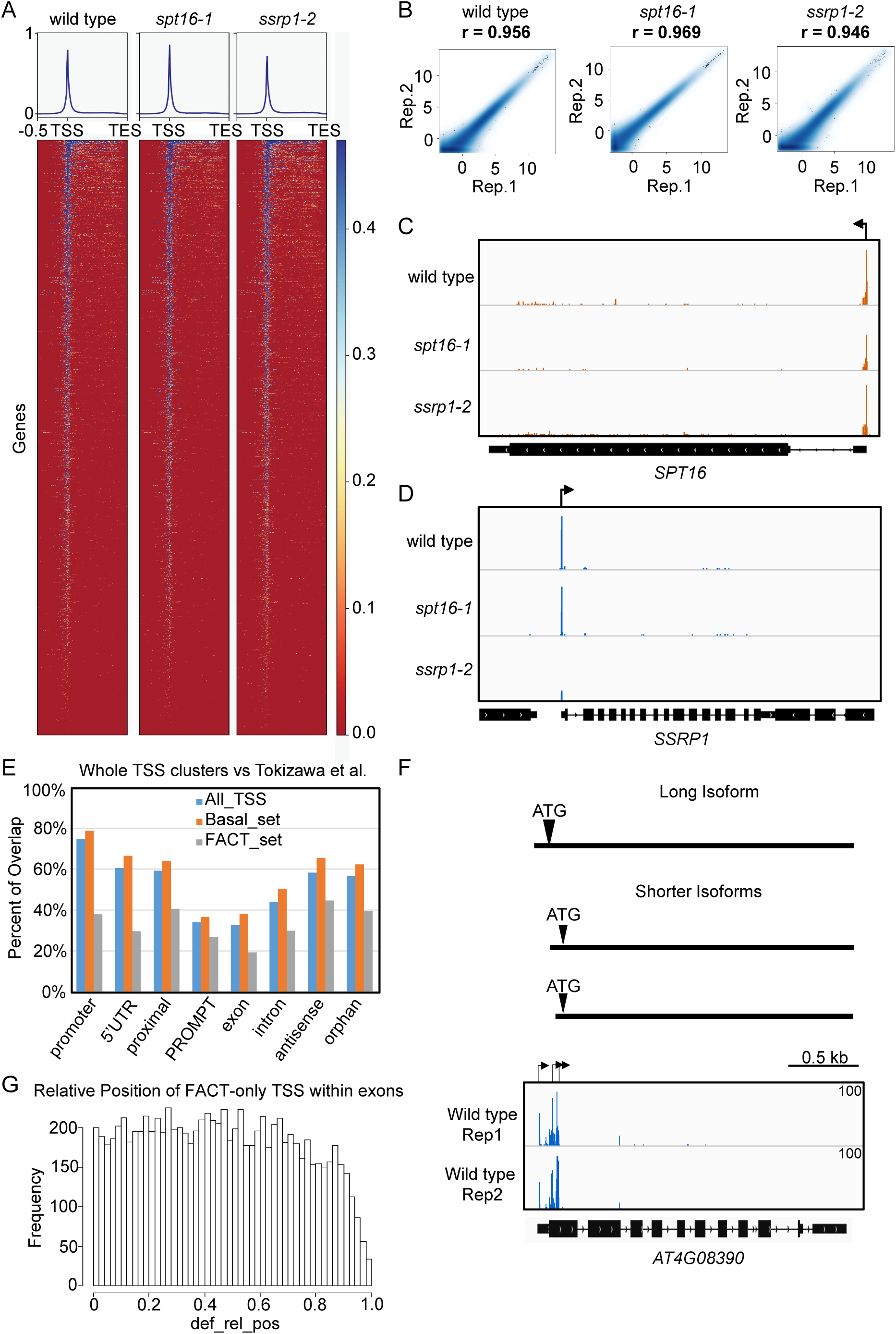
**Genome-wide TSS mapping in *Arabidopsis***. (A) TSS-seq read distribution across expressed *Arabidopsis* genes from −0.5 kb from transcription start site (TSS) to transcription end site (TES) in wild type, *spt16-1*, and *ssrp1-2*. (B) Reproducibility of two TSS-seq replicates (Rep.) in wild type, *spt16-1*, and *ssrp1-2*. (C) Screenshot of reduced TSS peak at *SPT16* gene in *spt16-1* mutant. (D) Screenshot of reduced TSS peak at *SSRP1* gene in *ssrp1-2* mutant. (E) The fraction of TSS clusters which overlap with CAGE peak summits reported by Tokizawa *et al*., 2017. (F) Screenshot of different TSSs corresponding to alternative mRNA isoforms of the *At4G08390* gene. The shorter isoforms utilize a second in-frame ATG to produce an N-terminally truncated protein that is differentially targeted within the cell (Obara et al., 2002). (G) Distribution of FACT-specific TSS along exons.

## References

Antosz, W., Pfab, A., Ehrnsberger, H.F., Holzinger, P., Kollen, K., Mortensen, S.A., Bruckmann, A., Schubert, T., Langst, G., Griesenbeck, J., et al. (2017). The Composition of the Arabidopsis Rna Polymerase Ii Transcript Elongation Complex Reveals the Interplay between Elongation and Mrna Processing Factors. Plant Cell 29, 854–870.

Ard, R., and Allshire, R.C. (2016). Transcription-Coupled Changes to Chromatin Underpin Gene Silencing by Transcriptional Interference. Nucleic Acids Res 44, 10619–10630.

Ard, R., Allshire, R.C., and Marquardt, S. (2017). Emerging Properties and Functional Consequences of Noncoding Transcription. Genetics 207, 357–367.

Arner, E., Daub, C.O., Vitting-Seerup, K., Andersson, R., Lilje, B., Drablos, F., Lennartsson, A., Ronnerblad, M., Hrydziuszko, O., Vitezic, M., et al. (2015). Transcribed Enhancers Lead Waves of Coordinated Transcription in Transitioning Mammalian Cells. Science 347, 1010–1014.

Bannister, A.J., Schneider, R., Myers, F.A., Thorne, A.W., Crane-Robinson, C., and Kouzarides, T. (2005). Spatial Distribution of Di-and Tri-Methyl Lysine 36 of Histone H3 at Active Genes. J Biol Chem 280, 17732–17736.

Bell, O., Wirbelauer, C., Hild, M., Scharf, A.N., Schwaiger, M., MacAlpine, D.M., Zilbermann, F., van Leeuwen, F., Bell, S.P., Imhof, A., et al. (2007). Localized H3k36 Methylation States Define Histone H4k16 Acetylation During Transcriptional Elongation in Drosophila. EMBO J 26, 4974–4984.

Belotserkovskaya, R., Oh, S., Bondarenko, V.A., Orphanides, G., Studitsky, V.M., and Reinberg, D. (2003). Fact Facilitates Transcription-Dependent Nucleosome Alteration. Science 301, 1090–1093.

Bouton, S., Leboeuf, E., Mouille, G., Leydecker, M.T., Talbotec, J., Granier, F., Lahaye, M., Hofte, H., and Truong, H.N. (2002). Quasimodo1 Encodes a Putative Membrane-Bound Glycosyltransferase Required for Normal Pectin Synthesis and Cell Adhesion in Arabidopsis. Plant Cell 14, 2577–2590.

Buratowski, S. (2009). Progression through the Rna Polymerase Ii Ctd Cycle. Mol Cell 36, 541–546.

Carrozza, M.J., Li, B., Florens, L., Suganuma, T., Swanson, S.K., Lee, K.K., Shia, W.J., Anderson, S., Yates, J., Washburn, M.P., et al. (2005). Histone H3 Methylation by Set2 Directs Deacetylation of Coding Regions by Rpd3s to Suppress Spurious Intragenic Transcription. Cell 123, 581–592.

Carvalho, S., Raposo, A.C., Martins, F.B., Grosso, A.R., Sridhara, S.C., Rino, J., Carmo-Fonseca, M., and de Almeida, S.F. (2013). Histone Methyltransferase Setd2 Coordinates Fact Recruitment with Nucleosome Dynamics During Transcription. Nucleic Acids Res 41, 2881–2893.

Cheung, V., Chua, G., Batada, N.N., Landry, C.R., Michnick, S.W., Hughes, T.R., and Winston, F. (2008). Chromatin-and Transcription-Related Factors Repress Transcription from within Coding Regions Throughout the Saccharomyces Cerevisiae Genome. PLoS Biol 6, e277.

Clark-Adams, C.D., and Winston, F. (1987). The Spt6 Gene Is Essential for Growth and Is Required for Delta-Mediated Transcription in Saccharomyces Cerevisiae. Mol Cell Biol 7, 679–686.

Clough, S.J., and Bent, A.F. (1998). Floral Dip: A Simplified Method for Agrobacterium-Mediated Transformation of Arabidopsis Thaliana. Plant J 16, 735–743.

Corden, J.L. (2013). Rna Polymerase Ii C-Terminal Domain: Tethering Transcription to Transcript and Template. Chem Rev 113, 8423–8455.

Davuluri, R.V., Suzuki, Y., Sugano, S., Plass, C., and Huang, T.H. (2008). The Functional Consequences of Alternative Promoter Use in Mammalian Genomes. Trends Genet 24, 167–177.

Descostes, N., Heidemann, M., Spinelli, L., Schuller, R., Maqbool, M.A., Fenouil, R., Koch, F., Innocenti, C., Gut, M., Gut, I., et al. (2014). Tyrosine Phosphorylation of Rna Polymerase Ii Ctd Is Associated with Antisense Promoter Transcription and Active Enhancers in Mammalian Cells. Elife 3, e02105.

Dobin, A., Davis, C.A., Schlesinger, F., Drenkow, J., Zaleski, C., Jha, S., Batut, P., Chaisson, M., and Gingeras, T.R. (2013). Star: Ultrafast Universal Rna-Seq Aligner. Bioinformatics 29, 15–21.

Dorighi, K.M., Swigut, T., Henriques, T., Bhanu, N.V., Scruggs, B.S., Nady, N., Still, C.D., 2nd, Garcia, B.A., Adelman, K., and Wysocka, J. (2017). Mll3 and Mll4 Facilitate Enhancer Rna Synthesis and Transcription from Promoters Independently of H3k4 Monomethylation. Mol Cell 66, 568–576 e564.

Duroux, M., Houben, A., Ruzicka, K., Friml, J., and Grasser, K.D. (2004). The Chromatin Remodelling Complex Fact Associates with Actively Transcribed Regions of the Arabidopsis Genome. Plant J 40, 660–671.

Earley, K.W., Haag, J.R., Pontes, O., Opper, K., Juehne, T., Song, K., and Pikaard, C.S. (2006). Gateway-Compatible Vectors for Plant Functional Genomics and Proteomics. Plant J 45, 616–629.

Eick, D., and Geyer, M. (2013). The Rna Polymerase Ii Carboxy-Terminal Domain (Ctd) Code. Chem Rev 113, 8456–8490.

Feng, J., Gan, H., Eaton, M.L., Zhou, H., Li, S., Belsky, J.A., MacAlpine, D.M., Zhang, Z., and Li, Q. (2016). Noncoding Transcription Is a Driving Force for Nucleosome Instability in Spt16 Mutant Cells. Mol Cell Biol 36, 1856–1867.

Francia, S., Michelini, F., Saxena, A., Tang, D., de Hoon, M., Anelli, V., Mione, M., Carninci, P., and d’Adda di Fagagna, F. (2012). Site-Specific Dicer and Drosha Rna Products Control the DNA-Damage Response. Nature 488, 231–235.

Gerstein, M.B., Lu, Z.J., Van Nostrand, E.L., Cheng, C., Arshinoff, B.I., Liu, T., Yip, K.Y., Robilotto, R., Rechtsteiner, A., Ikegami, K., et al. (2010). Integrative Analysis of the Caenorhabditis Elegans Genome by the Modencode Project. Science 330, 1775–1787.

Grini, P.E., Thorstensen, T., Alm, V., Vizcay-Barrena, G., Windju, S.S., Jorstad, T.S., Wilson, Z.A., and Aalen, R.B. (2009). The Ash1 Homolog 2 (Ashh2) Histone H3 Methyltransferase Is Required for Ovule and Anther Development in Arabidopsis. PLoS One 4, e7817.

Guenther, M.G., Levine, S.S., Boyer, L.A., Jaenisch, R., and Young, R.A. (2007). A Chromatin Landmark and Transcription Initiation at Most Promoters in Human Cells. Cell 130, 77–88.

Hainer, S.J., Charsar, B.A., Cohen, S.B., and Martens, J.A. (2012). Identification of Mutant Versions of the Spt16 Histone Chaperone That Are Defective for Transcription-Coupled Nucleosome Occupancy in Saccharomyces Cerevisiae. G3 (Bethesda) 2, 555–567.

Hajheidari, M., Koncz, C., and Eick, D. (2013). Emerging Roles for Rna Polymerase Ii Ctd in Arabidopsis. Trends Plant Sci 18, 633–643.

Hedtke, B., and Grimm, B. (2009). Silencing of a Plant Gene by Transcriptional Interference. Nucleic Acids Res 37, 3739–3746.

Hondele, M., and Ladurner, A.G. (2013). Catch Me If You Can: How the Histone Chaperone Fact Capitalizes on Nucleosome Breathing. Nucleus 4, 443–449.

Ikeda, Y., Kinoshita, Y., Susaki, D., Ikeda, Y., Iwano, M., Takayama, S., Higashiyama, T., Kakutani, T., and Kinoshita, T. (2011). Hmg Domain Containing Ssrp1 Is Required for DNA Demethylation and Genomic Imprinting in Arabidopsis. Dev Cell 21, 589–596.

Inagaki, S., Takahashi, M., Hosaka, A., Ito, T., Toyoda, A., Fujiyama, A., Tarutani, Y., and Kakutani, T. (2017). Gene-Body Chromatin Modification Dynamics Mediate Epigenome Differentiation in Arabidopsis. EMBO J 36, 970–980.

Jamai, A., Puglisi, A., and Strubin, M. (2009). Histone Chaperone Spt16 Promotes Redeposition of the Original H3-H4 Histones Evicted by Elongating Rna Polymerase. Mol Cell 35, 377–383.

Jefferson, R.A., Kavanagh, T.A., and Bevan, M.W. (1987). Gus Fusions - Beta-Glucuronidase as a Sensitive and Versatile Gene Fusion Marker in Higher-Plants. Embo Journal 6, 3901–3907.

Jensen, T.H., Jacquier, A., and Libri, D. (2013). Dealing with Pervasive Transcription. Mol Cell 52, 473–484.

Jeronimo, C., Watanabe, S., Kaplan, C.D., Peterson, C.L., and Robert, F. (2015). The Histone Chaperones Fact and Spt6 Restrict H2a.Z from Intragenic Locations. Mol Cell 58, 1113–1123.

Kaplan, C.D., Laprade, L., and Winston, F. (2003). Transcription Elongation Factors Repress Transcription Initiation from Cryptic Sites. Science 301, 1096–1099.

Keogh, M.C., Kurdistani, S.K., Morris, S.A., Ahn, S.H., Podolny, V., Collins, S.R., Schuldiner, M., Chin, K., Punna, T., Thompson, N.J., et al. (2005). Cotranscriptional Set2 Methylation of Histone H3 Lysine 36 Recruits a Repressive Rpd3 Complex. Cell 123, 593–605.

Kharchenko, P.V., Alekseyenko, A.A., Schwartz, Y.B., Minoda, A., Riddle, N.C., Ernst, J., Sabo, P.J., Larschan, E., Gorchakov, A.A., Gu, T., et al. (2011). Comprehensive Analysis of the Chromatin Landscape in Drosophila Melanogaster. Nature 471, 480–485.

Lawrence, M., Huber, W., Pages, H., Aboyoun, P., Carlson, M., Gentleman, R., Morgan, M.T., and Carey, V.J. (2013). Software for Computing and Annotating Genomic Ranges. PLoS Comput Biol 9, e1003118.

Li, B., Carey, M., and Workman, J.L. (2007). The Role of Chromatin During Transcription. Cell 128, 707–719.

Li, H., Handsaker, B., Wysoker, A., Fennell, T., Ruan, J., Homer, N., Marth, G., Abecasis, G., Durbin, R., and Genome Project Data Processing, S. (2009). The Sequence Alignment/Map Format and Samtools. Bioinformatics 25, 2078–2079.

Lloyd, J., and Meinke, D. (2012). A Comprehensive Dataset of Genes with a Loss-of-Function Mutant Phenotype in Arabidopsis. Plant Physiol 158, 1115–1129.

Lolas, I.B., Himanen, K., Gronlund, J.T., Lynggaard, C., Houben, A., Melzer, M., Van Lijsebettens, M., and Grasser, K.D. (2010). The Transcript Elongation Factor Fact Affects Arabidopsis Vegetative and Reproductive Development and Genetically Interacts with Hub1/2. Plant J 61, 686–697.

Mahrez, W., Arellano, M.S., Moreno-Romero, J., Nakamura, M., Shu, H., Nanni, P., Kohler, C., Gruissem, W., and Hennig, L. (2016). H3k36ac Is an Evolutionary Conserved Plant Histone Modification That Marks Active Genes. Plant Physiol 170, 1566–1577.

Malone, E.A., Clark, C.D., Chiang, A., and Winston, F. (1991). Mutations in Spt16/Cdc68 Suppress Cis-and Trans-Acting Mutations That Affect Promoter Function in Saccharomyces Cerevisiae. Mol Cell Biol 11, 5710–5717.

Marquardt, S., Hazelbaker, D.Z., and Buratowski, S. (2011). Distinct Rna Degradation Pathways and 3’ Extensions of Yeast Non-Coding Rna Species. Transcription 2, 145–154.

Marquardt, S., Raitskin, O., Wu, Z., Liu, F., Sun, Q., and Dean, C. (2014). Functional Consequences of Splicing of the Antisense Transcript Coolair on Flc Transcription. Mol Cell 54, 156–165.

Mayer, A., Lidschreiber, M., Siebert, M., Leike, K., Soding, J., and Cramer, P. (2010). Uniform Transitions of the General Rna Polymerase Ii Transcription Complex. Nat Struct Mol Biol 17, 272–1278.

Mellor, J., Woloszczuk, R., and Howe, F.S. (2016). The Interleaved Genome. Trends Genet 32, 57–71.

Nakagawa, T., Kurose, T., Hino, T., Tanaka, K., Kawamukai, M., Niwa, Y., Toyooka, K., Matsuoka, K., Jinbo, T., and Kimura, T. (2007). Development of Series of Gateway Binary Vectors, Pgwbs, for Realizing Efficient Construction of Fusion Genes for Plant Transformation. J Biosci Bioeng 104, 34–41.

Neri, F., Rapelli, S., Krepelova, A., Incarnato, D., Parlato, C., Basile, G., Maldotti, M., Anselmi, F., and Oliviero, S. (2017). Intragenic DNA Methylation Prevents Spurious Transcription Initiation. Nature 543, 72–77.

Ng, D.W., Wang, T., Chandrasekharan, M.B., Aramayo, R., Kertbundit, S., and Hall, T.C. (2007). Plant Set Domain-Containing Proteins: Structure, Function and Regulation. Biochim Biophys Acta 1769, 316–329.

Ntini, E., Jarvelin, A.I., Bornholdt, J., Chen, Y., Boyd, M., Jorgensen, M., Andersson, R., Hoof, I., Schein, A., Andersen, P.R., et al. (2013). Polyadenylation Site-Induced Decay of Upstream Transcripts Enforces Promoter Directionality. Nat Struct Mol Biol 20, 923–928.

Obara, K., Sumi, K., and Fukuda, H. (2002). The Use of Multiple Transcription Starts Causes the Dual Targeting of Arabidopsis Putative Monodehydroascorbate Reductase to Both Mitochondria and Chloroplasts. Plant Cell Physiol 43, 697–705.

Orphanides, G., Wu, W.H., Lane, W.S., Hampsey, M., and Reinberg, D. (1999). The Chromatin-Specific Transcription Elongation Factor Fact Comprises Human Spt16 and Ssrp1 Proteins. Nature 400, 284–288.

Parsley, K., and Hibberd, J.M. (2006). The Arabidopsis Ppdk Gene Is Transcribed from Two Promoters to Produce Differentially Expressed Transcripts Responsible for Cytosolic and Plastidic Proteins. Plant Mol Biol 62, 339–349.

Pelechano, V., Wei, W., and Steinmetz, L.M. (2016). Genome-Wide Quantification of 5’-Phosphorylated Mrna Degradation Intermediates for Analysis of Ribosome Dynamics. Nat Protoc 11, 359–376.

Pokholok, D.K., Harbison, C.T., Levine, S., Cole, M., Hannett, N.M., Lee, T.I., Bell, G.W., Walker, K., Rolfe, P.A., Herbolsheimer, E., et al. (2005). Genome-Wide Map of Nucleosome Acetylation and Methylation in Yeast. Cell 122, 517–527.

Porrua, O., and Libri, D. (2015). Transcription Termination and the Control of the Transcriptome: Why, Where and How to Stop. Nat Rev Mol Cell Biol 16, 190–202.

Price, B.D., and D’Andrea, A.D. (2013). Chromatin Remodeling at DNA Double-Strand Breaks.152, 1344–1354.

Proudfoot, N.J. (1986). Transcriptional Interference and Termination between Duplicated Alpha-Globin Gene Constructs Suggests a Novel Mechanism for Gene Regulation. Nature 322, 562–565.

Proudfoot, N.J. (2016). Transcriptional Termination in Mammals: Stopping the Rna Polymerase Ii Juggernaut. Science 352, aad9926.

Quinlan, A.R., and Hall, I.M. (2010). Bedtools: A Flexible Suite of Utilities for Comparing Genomic Features. Bioinformatics 26, 841–842.

Roudier, F., Ahmed, I., Berard, C., Sarazin, A., Mary-Huard, T., Cortijo, S., Bouyer, D., Caillieux, E., Duvernois-Berthet, E., Al-Shikhley, L., et al. (2011). Integrative Epigenomic Mapping Defines Four Main Chromatin States in Arabidopsis. Embo Journal 30, 1928–1938.

Schwabish, M.A., and Struhl, K. (2004). Evidence for Eviction and Rapid Deposition of Histones Upon Transcriptional Elongation by Rna Polymerase Ii. Mol Cell Biol 24, 10111–10117.

Tokizawa, M., Kusunoki, K., Koyama, H., Kurotani, A., Sakurai, T., Suzuki, Y., Sakamoto, T., Kurata, T., and Yamamoto, Y.Y. (2017). Identification of Arabidopsis Genic and Non-Genic Promoters by Paired-End Sequencing of Tss Tags. Plant J 90, 587–605.

Ushijima, T., Hanada, K., Gotoh, E., Yamori, W., Kodama, Y., Tanaka, H., Kusano, M., Fukushima, A., Tokizawa, M., Yamamoto, Y.Y., et al. (2017). Light Controls Protein Localization through Phytochrome-Mediated Alternative Promoter Selection. Cell.

van Bakel, H., Tsui, K., Gebbia, M., Mnaimneh, S., Hughes, T.R., and Nislow, C. (2013). A Compendium of Nucleosome and Transcript Profiles Reveals Determinants of Chromatin Architecture and Transcription. PLoS Genet 9, e1003479.

Van Lijsebettens, M., and Grasser, K.D. (2014). Transcript Elongation Factors: Shaping Transcriptomes after Transcript Initiation. Trends Plant Sci 19, 717–726.

Venkatesh, S., Smolle, M., Li, H., Gogol, M.M., Saint, M., Kumar, S., Natarajan, K., and Workman, J.L. (2012). Set2 Methylation of Histone H3 Lysine 36 Suppresses Histone Exchange on Transcribed Genes. Nature 489, 452–455.

von Arnim, A.G., Jia, Q., and Vaughn, J.N. (2014). Regulation of Plant Translation by Upstream Open Reading Frames. Plant Sci 214, 1–12.

Wiesner, T., Lee, W., Obenauf, A.C., Ran, L., Murali, R., Zhang, Q.F., Wong, E.W., Hu, W., Scott, S.N., Shah, R.H., et al. (2015). Alternative Transcription Initiation Leads to Expression of a Novel Alk Isoform in Cancer. Nature 526, 453–457.

Wilson, M.D., Harreman, M., Taschner, M., Reid, J., Walker, J., Erdjument-Bromage, H., Tempst, P., and Svejstrup, J.Q. (2013). Proteasome-Mediated Processing of Def1, a Critical Step in the Cellular Response to Transcription Stress. Cell 154, 983–995.

Zhang, F., Qi, B., Wang, L., Zhao, B., Rode, S., Riggan, N.D., Ecker, J.R., and Qiao, H. (2016a). Ein2-Dependent Regulation of Acetylation of Histone H3k14 and Non-Canonical Histone H3k23 in Ethylene Signalling. Nat Commun 7, 13018.

Zhang, T., Marand, A.P., and Jiang, J. (2016b). Plantdhs: A Database for Dnase I Hypersensitive Sites in Plants. Nucleic Acids Res 44, D1148–1153.

Zhao, Z., Yu, Y., Meyer, D., Wu, C., and Shen, W.H. (2005). Prevention of Early Flowering by Expression of Flowering Locus C Requires Methylation of Histone H3 K36. Nat Cell Biol 7, 1256–1260.

Zipfel, C., Kunze, G., Chinchilla, D., Caniard, A., Jones, J.D., Boller, T., and Felix, G. (2006). Perception of the Bacterial Pamp Ef-Tu by the Receptor Efr Restricts Agrobacterium-Mediated Transformation. Cell 125, 749–760.

